# Kinetic models for PET displacement studies

**DOI:** 10.1101/2022.11.25.517914

**Authors:** Gjertrud Louise Laurell, Pontus Plavén-Sigray, Annette Johansen, Nakul Ravi Raval, Arafat Nasser, Clara Aabye Madsen, Jacob Madsen, Hanne Demant Hansen, Lene Lundgaard Donovan, Gitte Moos Knudsen, Adriaan A Lammertsma, R Todd Ogden, Claus Svarer, Martin Schain

## Abstract

The traditional design of PET target engagement studies is based on a baseline scan and one or more scans after drug administration. We here evaluate an alternative design in which the drug is administered during an on-going scan (i.e., a displacement study). This approach results both in lower radiation exposure and lower costs. Existing kinetic models assume steady state. This condition is not present during a drug displacement and consequently, our aim here was to develop kinetic models for analysing PET displacement data.

We modified existing compartment models to accommodate a time-variant increase in occupancy following the pharmacological in-scan intervention. Since this implies the use of differential equations that cannot be solved analytically, we developed instead one approximate and one numerical solution. Through simulations, we show that if the occupancy is relatively high, it can be estimated without bias and with good accuracy. The models were applied to PET data from six pigs where [^11^C]UCB-J was displaced by intravenous brivaracetam. The dose-occupancy relationship estimated from these scans showed good agreement with occupancies calculated with Lassen plot applied to baseline-block scans of two pigs. In summary, the proposed models provide a framework to determine target occupancy from a single displacement scan.

## 1. Introduction

The discovery and development of drugs for the treatment of brain disorders is a challenging process requiring extensive resources, long timelines and significant investments.^1^ Over the past decades, imaging with positron emission tomography (PET) has become a valuable tool in CNS drug development. PET imaging with appropriate radioligands makes it possible to determine at an early stage whether a candidate drug penetrates the blood-brain barrier and binds to the target of interest *in vivo*. This helps to ensure that only suitable candidates will be advanced to subsequent trial phases, saving substantial resources.^2^

In the traditional experimental set-up for determination of target occupancy, two (or more) PET scans are acquired in each subject using a radiotracer that binds to the same target as the drug. Usually, one scan is acquired at baseline (i.e., without drug), and subsequent scan(s) after administration of the drug. The difference between outcome measures from baseline and follow-up scans is then used to determine occupancy, i.e., the fraction of targets occupied by the drug.^3^

The analysis of data from a PET occupancy study typically follows a three-step approach. First, a mathematical model (either based on arterial blood samples or a reference region) is used to quantify radiotracer uptake for each scan.^4^ Second, outcome measures from the different scans are combined to estimate the occupancy at the time of the post-drug scan.^3,5^ Last, all subjects are pooled in a occupancy plot (sometimes referred to as the *E*_*max*_ model) where the administered doses or plasma concentrations of the drug are related to the measured occupancies. This final step provides information of the drug’s affinity to the target, defined as the half maximum inhibitory concentration, *IC*_*50*_.

This established methodology leaves room for improvement. First, relying on multiple PET measurements may introduce unwanted variance in the data. It is often difficult to design the experiment in such a way that the intervention is the only difference between the scans, as other (e.g., time-related) factors may affect radiotracer uptake.^6–9^ Second, the test person is exposed to multiple doses of ionizing radiation. Third, PET is a relatively expensive research tool, so a method that requires two scans for each data point places an unnecessary burden on the research budget.

An alternative approach to determine drug occupancy is to administer the drug during an on-going PET scan and based on a single injection of radioligand. When the drug is administered, competitive binding causes displacement of the radiotracer. Deriving occupancy from such a study would result in reduced costs and lower radiation exposure, as each subject would need to undergo only one scan.

Unfortunately, standard pharmacokinetic models, routinely used to analyse PET data, cannot be applied to data obtained from a displacement scan, as these models rely on the assumption of steady-state throughout a scan.^10^ This assumption implies that all model parameters remain constant over time, i.e., the modelled system is assumed to be time-invariant. This assumption is violated when a competing drug is administered during an on-going scan. To quantify displacement studies, a new class of pharmacokinetic models needs to be developed to incorporate the perturbation of the steady state.

The idea to model competitive binding was already pioneered in the 1990s, when models to quantify release of endogenous dopamine were developed.^11–14^ Of the models relying on bolus injection, the only established model today is the neurotransmitter PET (ntPET), which has been used to evaluate dopamine release under various conditions.^15–18^ The ntPET, and variants thereof,^19–23^ rely on reference tissue models, i.e., quantification is performed using a reference region rather than an arterial input function. For many radiotracers, true reference regions do not exist for, due to ubiquitous expression of the target, although some degree of specific binding in the reference region may be tolerable in clinical studies. In pharmacological intervention studies, however, specific binding in the reference region is particularly problematic, as blocking in both target and reference region can result in a complicated bias in the occupancy estimates.

Here, we present a pharmacokinetic model that describes radiotracer kinetics during a displacement scan based on an arterial input function rather than a reference region. The model is based on one-tissue compartment model (1TCM) kinetics. The corresponding two-tissue compartment model (2TCM) is presented in section A of the supplementary material. Because the competing drug will perturb the system’s steady state, analytical solutions to the model equations do not exist. Instead, we present two alternative approaches: an approximate analytical solution that is derived by introducing assumptions on the accumulation of the competing drug in brain, and one numerical solution. The performance of the model and solutions are evaluated using simulations, and applied to pig [^11^C]UCB-J PET scans,^24^ with brivaracetam displacement.

## 2. Materials and methods

### 2.1 Theory

#### 2.1.1 The occupancy model

When radioligand displacement is induced by the introduction of a competing cold ligand (drug) it is assumed that the change is caused by reduction in specific radioligand binding only. This can be modelled by defining an occupancy function, *∂*(*t*), with 0 ≤ *∂*(*t*) ≤ 1, that acts on the concentration of available binding sites, *B*_*avail*_. We typically have little knowledge about the drug concentration time profile in brain tissue *in vivo*. To set *∂*(*t*), we defined a set of conditions to be fulfilled. The occupancy model should

1. be monotone non-decreasing
2. be continuous and differentiable in all time points (i.e., a smooth growth)
3. be 0 at the time of drug administration

To fulfil the conditions above, we modified a model originally developed in agricultural sciences to predict crop growth rates.^25^ Our model for the occupancy function *∂*(*t*) is

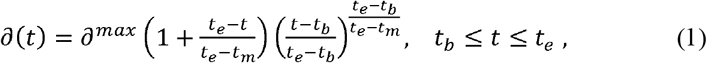

where *∂*^*max*^ is the maximal occupancy reached (0 ≤ *∂*^*max*^ ≤1), *t_e_* is the end time of the growth (i.e., the time at which *∂*^*max*^ is reached), *t*_*b*_ is the begin-time of the growth (i.e., the time of intervention), and *t*_*m*_ is the time during which *∂*′(*t*) reaches a maximum (i.e., *t*_*m*_ will control the steepness of *∂*(*t*)). The function *∂*(*t*) will show a sigmoidal growth within the interval *t*_*b*_ ≤ *t* ≤ *t*_*e*_, is exactly 0 at *t* = *t*_*b*_, and can allow asymmetric or symmetric growth curves depending on the choices of *t*_*e*_ and *t*_*m*_. Examples of *∂*(*t*) are shown in Figure 1.

**Figure 1:**
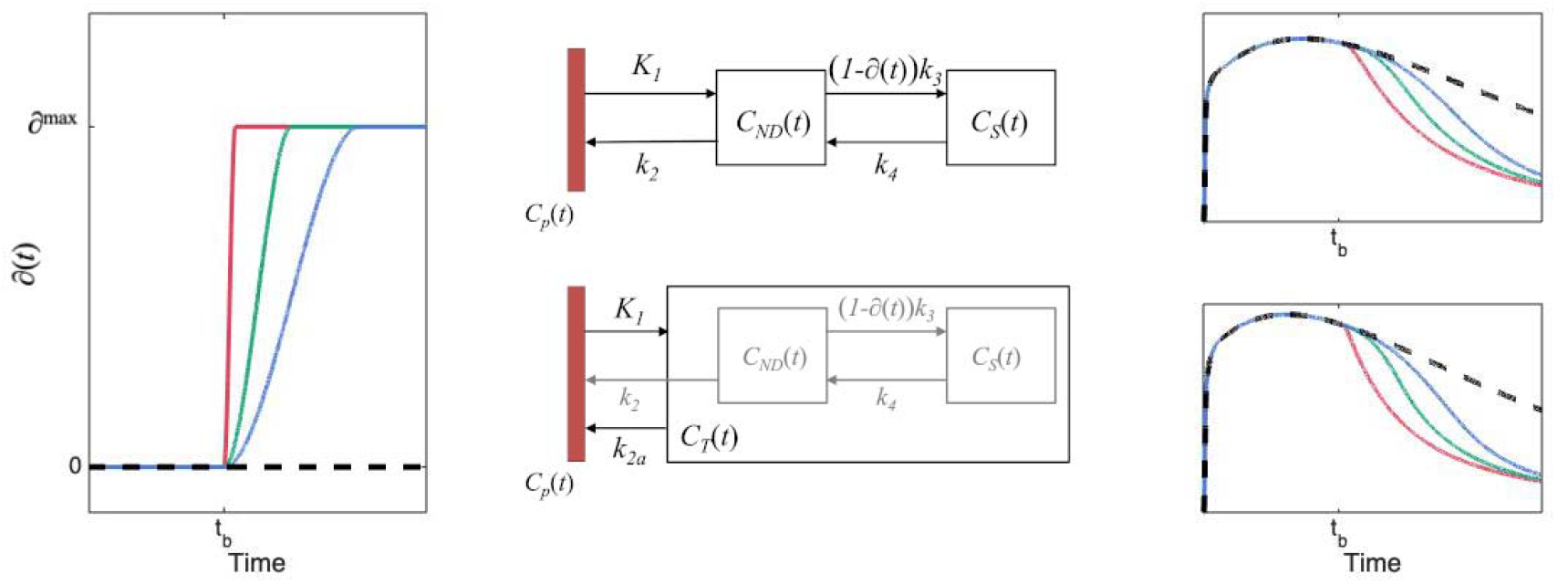
The left panel shows the occupancy function, *∂*(t) as defined in Eq.1, for 3 different choices of t_e_. The middle panels show schematic diagrams of the intervention models for 2TCM and 1TCM, respectively. The right panels show time-activity curves originating from each compartmental model using the choices for *∂*(t) depicted in the left panel.

#### 2.1.2 Displacement model

The 2TCM is the most common pharmacokinetic model used in quantification of brain PET data. In the 2TCM, the rate of exchange between compartments is determined by the constants *K*_1_, *k*_2_, *k*_3_ and *k*_4_ (Figure 1). The rate constant *k*_*3*_ is linearly dependent on the concentration of will have a negligible impact on available targets (*B*_*avail*_), *k*_3_ = *f*_*ND*_ *k*_*on*_ *B*_*avail*_.^3,4^ We assume that a reduction of available targets will have a negligible impact on both the association rate constant *k*_*on*_ and the fraction of free tracer in the non-displaceable compartment, *f*_*ND*_. It follows that a time dependent reduction of available targets, i.e., (1 − *∂*(*t*))*B*_*avail*_ will affect *k*_*3*_ equally, i.e., *f*_*ND*_ *k*_*on*_ · (1 − *∂*(*t*))*B*_*avail*_ = (1 − *∂*(*t*))*k*_3_. With this, the 2TCM can be modified to accommodate an increase in occupancy, starting at some time *t*_*b*_ after radiotracer injection (details and equations provided in section A of the supplementary material).

The pharmacokinetics of some radiotracers are reasonably well approximated by a 1TCM. In the 1TCM, the compartments corresponding to specific and non-displaceable uptake are collapsed into a single compartment, where rate constants *K*_*1*_ and *k*_*2*_ describe the transfer rate of radiotracer to and from that compartment. To modify the 1TCM to accommodate displacement, we adapted the framework of the simplified reference tissue model, where the compartments for specific and non-displaceable binding are presumed to latently reside within the model configuration (Figure 1).^26^ Setting equal the distribution volumes for the 1- and 2 tissue compartment configurations, a relationship between the apparent efflux rate constant, *k*_*2a*_, and *k*_*2*_-*k*_*4*_ from the latently present 2TCM configuration can be derived, 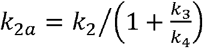. modification enables the introduction of an occupancy parameter to act on *k*_*3*_ in the 1TCM configuration. A schematic for the model is shown in Figure 1, and the corresponding differential equation becomes

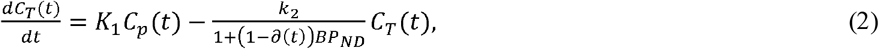

where *C*_*p*_ (*t*) is the metabolite corrected arterial plasma input function, *BP*_*ND*_ = *k*_3_/*k*_4_, and *∂*(*t*) as defined by

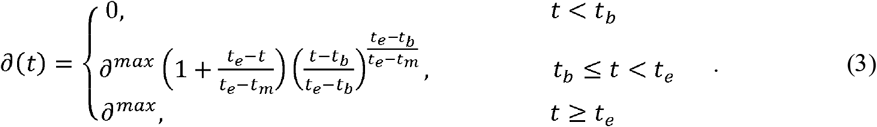

As indicated in equation (3), the competing drug is assumed not to wash out during the course of the scan (see Discussion).

#### 2.1.3 The single step solution

In contrast to the standard kinetic models, equations (2) describe a time-variant system, and common tools for finding analytical solutions are not strictly defined. To find solutions for the differential equations (2), we placed some restrictions on the occupancy function,.

For a drug acting rapidly on the target, i.e., quickly reaching the maximal occupancy attainable at the administered dose, we assume that takes the form of a step function, i.e.,

With this simplification, we can partition the PET time-activity curve (TAC) into two segments (i.e., before and after the time at which the drug is assumed to act on the system, *t*_*s*_) and apply the 1TCM separately to each segment. We assume that the rate constants are the same for the two segments, but the differential equation for the segment after the step (*t* > *t*_*s*_) will come with non-zero initial values. The initial values for the segment are set to the *t* > *t*_*s*_ values at the endpoint of the *t* > *t*_*s*_ segment.

Let 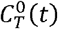 and 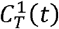 denote the tissue concentrations before and after administration of the competing drug. The equations describing 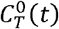 are the standard differential equations for the 1TCM (see equation 2, with *∂*(*t*) = 0). Setting τ = *t* − *t*_*s*_, the differential equation for *t* > *t*_*s*_ becomes

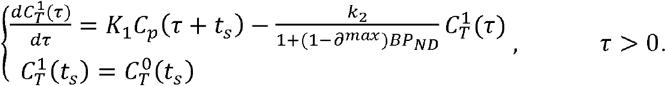

The solution to these differential equations becomes

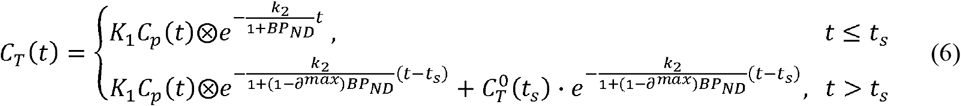

Fitting the single step solution thus means estimating a total of 4 model parameters: *K*_*1*_, *k*_*2*_ and *BP*_ND_ and *∂*^*max*^. In our implementation, we also included the fractional blood volume (*v*_*B*_), and the time at which the occupancy step occurs (*t*_*s*_), as free parameters. The reason to include *t*_*s*_ as a free parameter is to reduce errors caused by setting *∂*(*t*) to be a step function: by allowing the model to perform the step later than the time of intervention we submit that a better description of the data can be obtained. The equations for the single step solution of the 2TCM is provided in the supplementary material (section A.2).

#### 2.1.4 The Numerical Solution

The Euler Forward method was used to obtain a numerical solution to equation (2). Starting at known initial conditions, each next point on the model curves is calculated by,

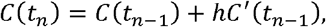

where *h* = *t*_*n*_ − *t*_*n*−1_ (i.e., linearity is assumed in the small time interval *t*_*n*−1_ < *t* < *t*_*n*_). Insertion of the model equations in (2) gives

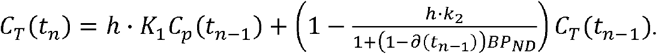

The occupancy function shown in equation 3 (Figure 1) was used for *∂*(*t*_*n*_), and *h* was set to 0.5 seconds. The corresponding numerical solution for the 2TCM is presented in the supplementary material (section A.3).

#### 2.1.4 Fitting of multiple regions simultaneously

As seen in equation (2), the free model parameters *k*_2_, *∂*(*t*), and *BP*_*ND*_ appear only as a ratio, and as such, each of these parameters are free to take any value as long as the term 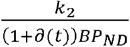 results in an adequate fit. Consequently, unless some constraints are placed in the optimization, neither of these parameters can be properly identified (see supplementary material, section B and supplementary figures 1 and 2).

In PET kinetic modelling, it is common to assume that the non-displaceable distribution volume (*V*_*ND*_) is the same across all included brain regions. In dose-occupancy studies, it is also common to assume that the fractional occupancy is the same across the brain. In fact, these two assumptions form the basis for many quantification strategies, including all reference tissue modelling as well as the Lassen Plot.^27^ We therefore constructed our optimizer so that *V*_*ND*_ and *∂*^*max*^ are shared across all included brain regions, whereas the other model parameters, i.e., *K*_1_, *BP*_*ND*_, and *ν*_*B*_, are free to vary across the brain. For the single step approach, the estimated time at which the model performs the jump, *t*_*s*_, was also treated as a global parameter. Similarly for the numerical solution, the estimated time at which the growth of the occupancy ends, *t*_*e*_, was estimated globally. For both methods, we used a nested approach to fit the models to the data. In an outer layer, the global parameters (*V*_*ND*_, *∂*^*max*^, *t*_*e*_/*t*_*s*_) were estimated with non-linear least squares. For each iteration of the outer layer, the remaining model parameters (*K*_*1*_, *V*_*ND*_, *v*_*B*_) were fitted for each ROI separately. More details about models and implementation can be found in the supplementary material (section C).

### 2.2 Simulations

#### 2.2.1 Generation of noise-free time activity curves

In simulations, we attempted to mimic the behaviour of [^11^C]UCB-J.^24^ For *C*_*p*_*(t)*, we used an arterial input function measured from a pig scan (baseline scan of experiment 1 in Table 1), where the measured activities after the peak were fitted with a tri-exponential function. The tissue rate constants were taken from table 1 in (Finnema et al., 2018)^28^, and set to result in *V*_*ND*_ = 4, and *V*_*T*_ ranging between 14.2 and 22.4. We simulated TACs for seven regions: putamen, temporal cortex, occipital cortex, frontal cortex, thalamus, cerebellum, and hippocampus.

**Table 1.** Overview of pig PET experiments.

| Experiment | Type | Dose [mg/kg] |
| --- | --- | --- |
| Experiment 1 | Baseline scan |  |
|  | Intervention scan | 0.1 |
|  | Block scan |  |
| Experiment 2 | Baseline scan |  |
|  | Intervention scan | 2 |
|  | Block scan |  |
| Experiment 3 | Intervention scan | 5 |
| Experiment 4 | Intervention scan | 1 |
| Experiment 5 | Intervention scan | 0.75 |
| Experiment 6 | Intervention scan | 0.2 |

To evaluate the performance of our methods, we simulated displacement TACs with two different types of drugs: one fast and one slower. In both cases, the time of injection of the drug was set at 60 minutes after radiotracer administration. For the fast-acting drug, the time *t*_*e*_ at which the drug reached it maximal occupancy was set to 65 minutes, for the slow-acting drug, *t*_*e*_ was set to 90 minutes. Scan durations were set to 150 minutes. For each of the drugs, we simulated displacement scans at three different occupancies (*∂*^*max*^): 25%, 50% and 75%. For each of the six different cases (combination of drug and occupancy) we simulated 1000 unique scans, with noise added as explained in *2*.*2*.*2 Generation of noise*. To generate the TACs we used the Euler Forward method and the differential equation for the 1TC displacement model (Equations 2 and 3). Some example TACs with different *t*_*e*_ are shown in Figure 1.

#### 2.2.2 Generation of noise

To create realistic noise, a previously proposed noise-model was used that allows for time-dependent variance.^29,30^

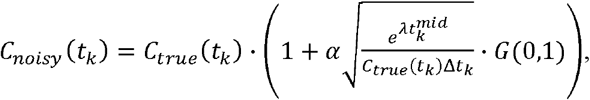

where *λ* is the decay constant for the isotope (in this case ^11^C), 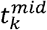 and Δ*t*_*k*_ are the mid-time and duration of frame *k*, respectively, and *G*(0,1) is a number sampled from a Gaussian distribution centered on 0 with a SD of 1. The scaling factor *α* was set to 5 in order to create noise on par with that of real experiments. Figures of example TACs at different noise levels can be found in the supplementary material (supplementary figures 3 and 4).

### 2.3 PET experiments

#### 2.3.1 Experimental procedure

[^11^C]UCB-J was largely synthesized as in Nabulsi et al., 2016,^24^ with some modifications.^31^ All animal experiments conformed to the European Commission’s Directive 2010/63/EU and the ARRIVE guidelines. The Danish Council of Animal Ethics had approved all procedures (Journal no. 2016-15-0201-01149). Six female domestic pigs (crossbreed of Yorkshire × Duroc × Landrace, mean weight 23.3 [range: 18-27] kg) were fully anesthetized and scanned using [^11^C]UCB-J administered as a bolus injection (injected dose: 441 [range: 344-528] MBq; injected mass: 0.69 [range: 0.08-2.77] μg). Two of the pigs (experiments 1 and 2) underwent three scans, i.e., baseline (120 min), displacement (150 min) and blocking (120 min) scans, on the same day. Brivaracetam (Briviact®, 10 mg/mL, UCB Pharma, Belgium) was administered i.v. during the displacement scan and served as a traditional blocking agent in the third scan, which was started approximately 120 min after the brivaracetam intervention. The remaining four pigs only underwent displacement scans. In all displacement scans, brivaracetam was administered i.v. over 20 seconds, 60 min after radioligand injection. A list of all experiments is shown in Table 1.

#### 2.3.3 PET data processing

PET scans were acquired on a High Resolution Research Tomograph (HRRT; Siemens, USA) and reconstructed using OP-3D-OSEM, including modelling of the point-spread function, with 16 subsets, 10 iterations and all standard corrections.^32^ Data was binned into the following time frames: 6 × 10, 6 × 20, 3 × 30, 9 × 60, 8 × 120, 4 × 180, 2 × 240, 1 × 360, 1 × 420, 1 × 600, 1 × 900 and 1 × 1680 s for the 120 min scans, and 6 × 10, 6 × 20, 6 × 30, 6 × 60, 4 × 120, 14 × 300, 8 × 150 s and 8 × 300 s for the 150 min scans. Attenuation correction was performed using the MAP-TR μ-map.^33^ Definition of brain regions of interests (ROIs) was performed using a dedicated pig brain template.^34^ The seven regions from the simulation experiment were also used here: putamen, temporal cortex, occipital cortex, frontal cortex, thalamus, cerebellum and hippocampus.

#### 2.3.4 Blood and plasma analyses

Radioactivity in arterial whole blood was measured continuously for the first 30 min of each scan using an Allogg ABSS autosampler (Allogg Technology, Sweden). Arterial blood was manually drawn at 3, 8, 10, 15, 30, 45, 59, 61, 65, 75, 90, 105, 120 and 150 min for measuring radioactivity in whole blood and plasma using a gamma counter (Cobra, 5003, Packard Instruments, Meriden, USA) that was cross-calibrated against the HRRT. Radio-HPLC was used to measure radioligand parent fractions.^35^ A more detailed account of the blood and plasma analyses can be found in the supplementary material (Section D).

In the baseline scan of experiment 2, (see Table 1), the parent fraction could not be estimated due to a technical failure. The parent fractions from the displacement and blocking scans conducted in the same animal were, however, very similar (absolute difference averaged across time was 6.0±4.1%, and difference in AUC was 2%). Therefore, for the baseline scan in experiment 2, the mean parent fraction from the corresponding displacement and blocking scans was used.

The concentration of brivaracetam in arterial plasma was analysed using UPLC-MS/MS (Filadelfia Epilepsy Hospital, Denmark). During the displacement scans seven blood samples (at approximately 1, 5, 15, 30, 45, 60 and 90 min after brivaracetam injection) were collected for this purpose, and during the block scans five samples (at approximately 3, 15, 45, 75 and 90 min after scan start) were collected.

## 3. Results

### 3.1 Simulation results

Figure 2 summarises results from the simulation experiment. It shows that occupancy estimation improves both with increasing drug speed and with increasing dose. The performance of the two methods (numerical solution and single-step approximation) were comparable throughout, especially for the higher occupancies, where the histograms are almost identical. While the occupancy estimates are approximately normally distributed for the higher occupancies, the distributions of estimates are slightly skewed for both methods at 25% occupancy. At this lower occupancy, there is also a bigger difference between the distributions, with the single-step solution, unlike the numerical solution, showing a slight tendency to overestimate *∂*^max^ (for the fast drug, median *∂*^max^ estimates were 25.2% with the numerical solution and 30.2% with the single step solution). Corresponding results for *V*_*ND*_ and *V*_*S*_ are found in supplementary figures 8 and 9, respectively.

**Figure 2:**
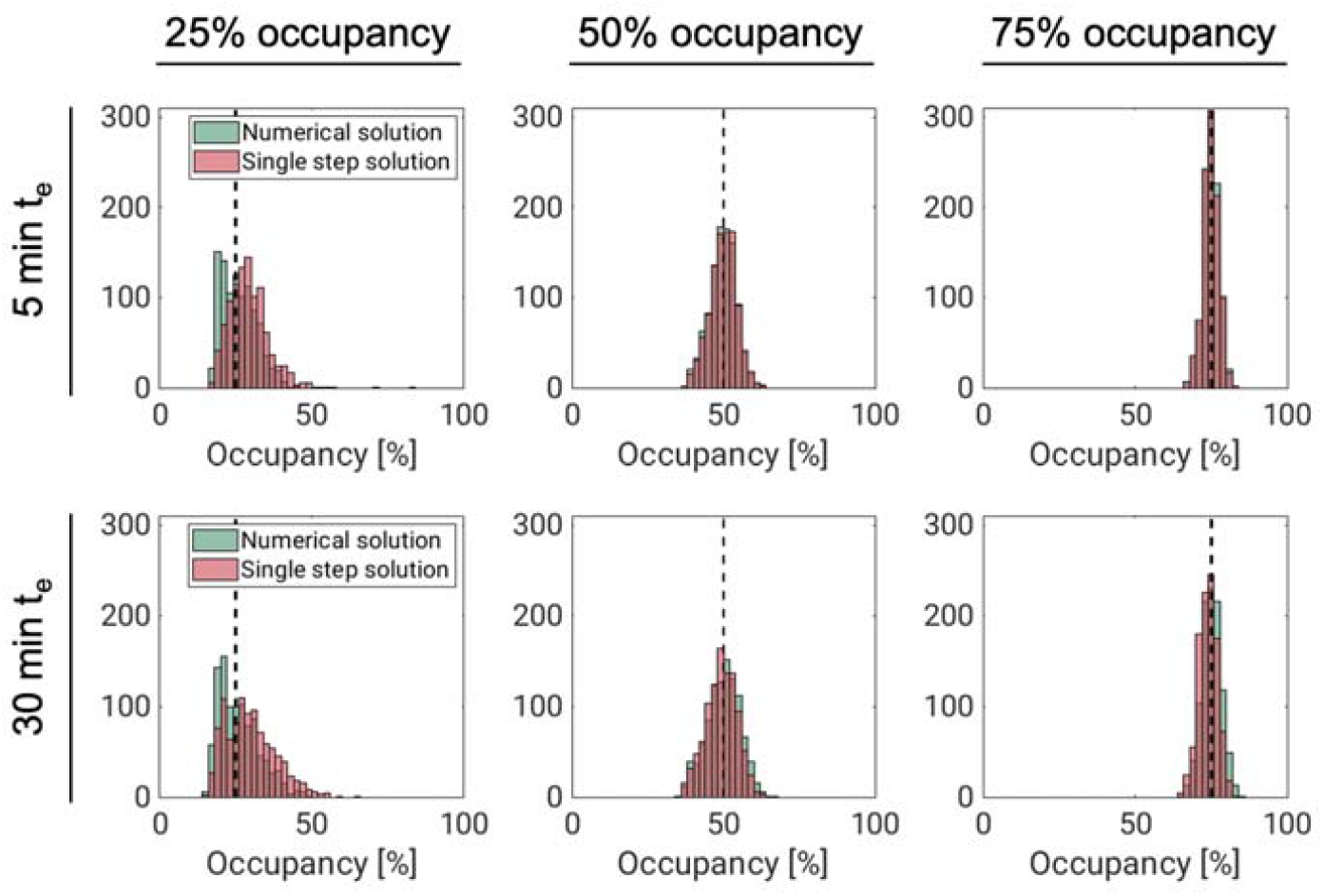
Results from simulating fast-(t_e_ = 5 min) and slow-acting (t_e_ = 30 min) drugs, displacing [^11^C]UCB-J binding. Each panel shows histograms of occupancy (*∂*^max^) estimates from the numerical solution (green) and single-step approximation (red). In each panel, the dashed black line corresponds to the true value for *∂*^max^.

### 3.2 Displacement models applied to real data

#### 3.2.1 Model fits

The 1TC displacement model was consistently able to describe the measured TACs using both the single-step approximation and the numerical solution. Figure 3 shows model fits to temporal cortex TACs, with both methods, for the largest and the lowest dose experiments. For the numerical solution, fits to all TACs, with residuals, for the same two scans can be found in supplementary figures 13 and 14, and normalized residuals for all six pig scans can be found in supplementary figure 15. The average ± SD total distribution volume in temporal cortex, across all displacement scans was 20.2 ± 3.7 mL/cm^3^ for both methods. The occupancies ranged from 41% to 86%.

**Figure 3:**
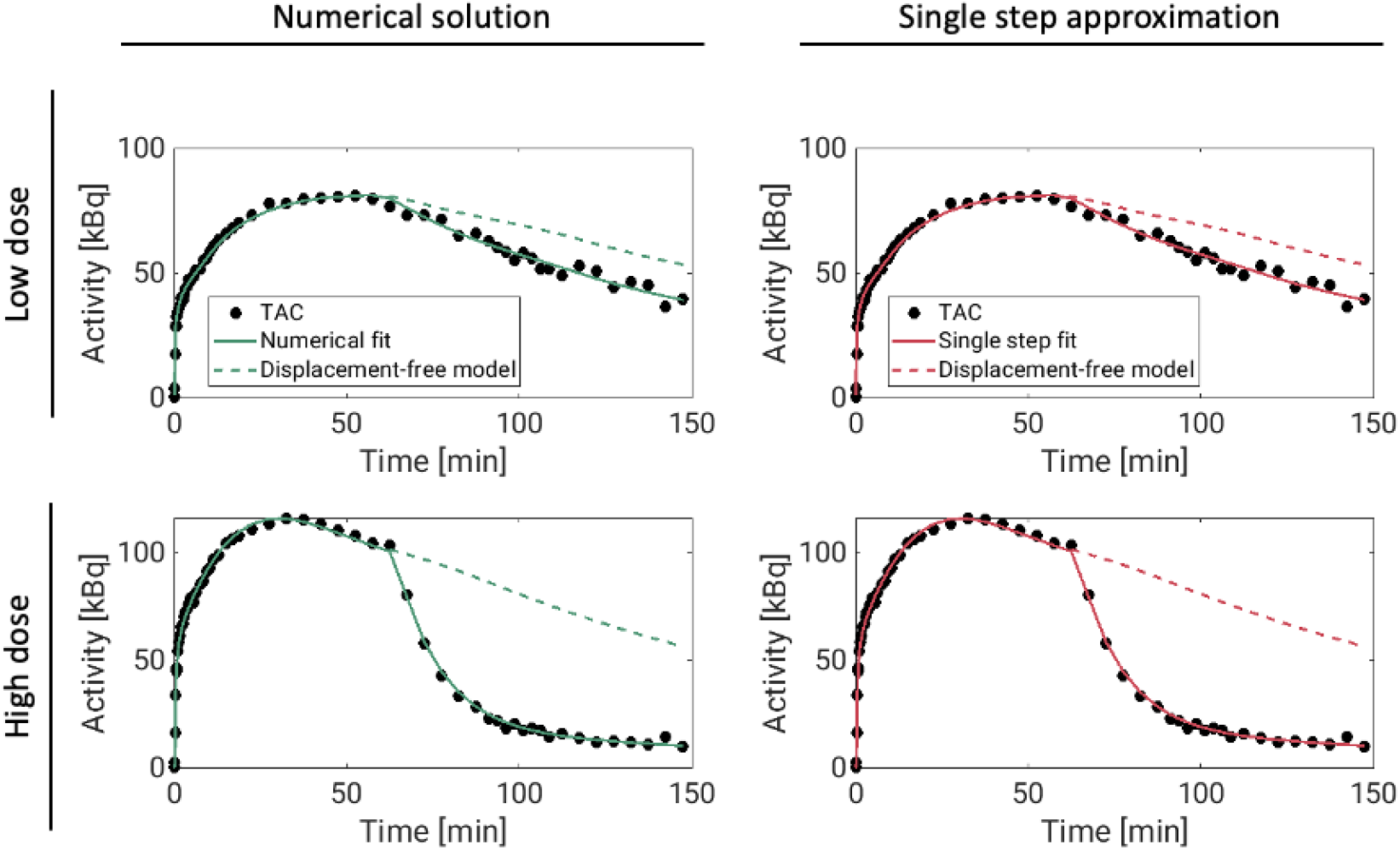
Displacement model fits (solid lines) to [^11^C]UCB-J temporal cortex TACs (dots) from two pig scans in which brivaracetam was administered i.v. 60 minutes after radiotracer injection. The dashed lines show model curves in the absence of displacement. These curves were generated from the estimated model parameters, with occupancy set to zero.

#### 3.2.2 Comparison with Lassen plots

Experiment 1 and 2 each had a baseline scan before and a blocking scan after the displacement scan. These scans were analysed using the traditional 1TCM, and occupancies were estimated using the Lassen plot.^5,27^ For both experiments, the Lassen occupancies in the block scans were lower than the estimates from the displacement scans (see figure 4, left panel). However, the plasma drug concentrations were also lower during the block scans, and the outcome of the Lassen-plot fit well with the dose-occupancy relationship estimated from the displacement scans (see dose-occupancy plots in figure 4). The Lassen-*V*_*ND*_s also showed good agreement with the ones calculated with the displacement model. For the high-dose pig scan, the Lassen plot returned a *V*_*ND*_ of 2.08, while the displacement model returned a *V*_*ND*_ of 1.85 with the numerical solution and 2.15 with the single step solution. For the low-dose pig scan, the Lassen-*V*_*ND*_ was 7.46, while the displacement model returned *V*_*ND*_ estimates of 7.47 with both solutions.

**Figure 4:**
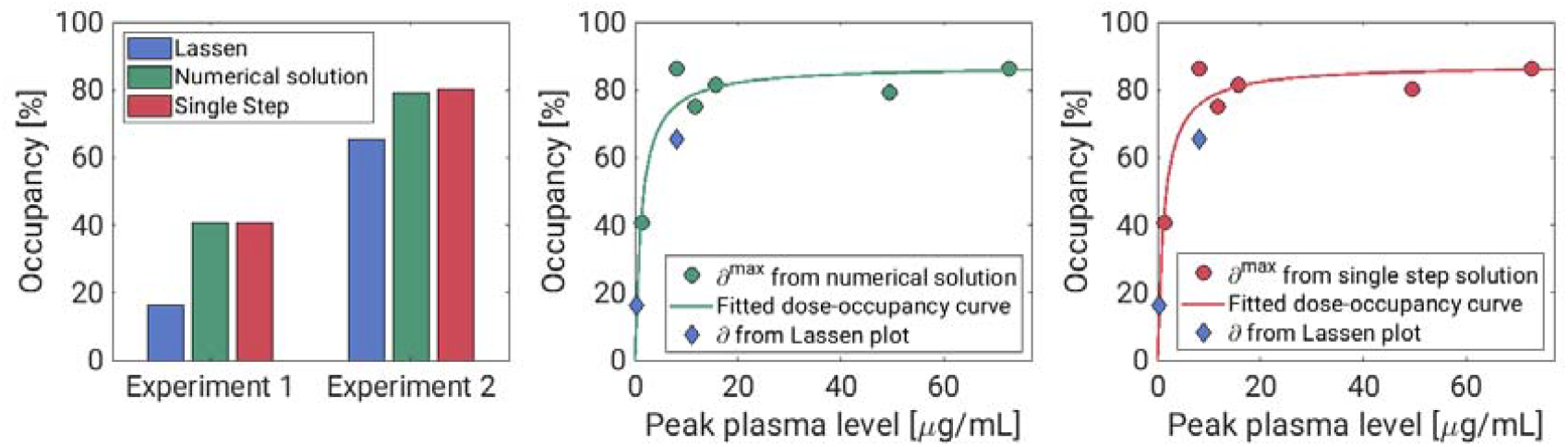
Results from pig experiment. The first tile show occupancy estimates for the two pigs that underwent both displacement and baseline-block scans. The Lassen plot occupancy estimates (from the block scan) are shown in blue, and the occupancy estimates from the displacement scans are shown in green (numerical solution) and red (single step solution). The two other tiles show the estimated occupancies plotted against peak plasma brivaracetam concentrations for the numerical solution and single step solutions, respectively. In both, the Lassen-occupancies plotted against the peak plasma value during the block scans are also included.

#### 3.2.3 Dose-occupancy model

Figure 4 shows occupancies (*∂*^*max*^) estimated from the displacement models for all pigs, plotted against the maximal plasma level of brivaracetam following its injection (*C*_*briva*_). These occupancies could be well described by the *E*_*max*_ model, *∂*^*max*^ = Δ_*max*_ *C*_*briva*_/ (C_*briva*_ + *IC*_50_), where *IC*_*50*_ is the drug’s half maximal inhibitory concentration, and Δ_*max*_ is the maximal attainable occupancy for the population. The estimated values for Δ_*max*_ were 87.3% for the numerical solution, and 87.6% for the single step approach. The corresponding values for *IC*_*50*_ were 1.26 *μ*g/mL and 1.27 *μ*g/mL for the numerical solution and single step, respectively.

## Discussion

In this study, we present a pharmacokinetic model capable of describing PET time-activity curves after a pharmacological intervention. We have developed a generic and flexible model that allows for increasing occupancy, and that is incorporated into the standard PET compartmental models, to describe a displacement of the radiotracer during the scan. Because the differential equations are time-variant, we present two new approaches for quantification of PET data with arterial input functions. In the single step solution, the effect of the drug intervention is approximated to be instantaneous, and the system can thus be assumed to be time-invariant both before and after the effect of the intervention occurs. The “extended simplified reference tissue model” (ESRTM) is based on the same idea, although it relies on a reference tissue rather than an arterial input function.^36^ The other solution is to solve the model differential equations using numerical methods. Here, we used the Euler Forward method together with a monotone, continuous and differentiable occupancy model.

The present results suggest that performing displacement scans is a viable alternative to the traditional two-scan setup to determine target engagement. We demonstrate the usefulness of the methods by displacing [^11^C]UCB-J with brivaracetam in pigs. Experiments 1 and 2 showed that the displacement model resulted in similar occupancy estimates as those obtained using the traditional Lassen plot. The estimated occupancies could be well described by the *E*_*max*_ model, which resulted in *IC*_*50*_ estimates of 1.26 *μ*g/mL for the single step method and 1.27 μg/mL for the numerical solution. The *E*_*max*_ model assumes that the drug concentration in plasma is constant but plasma brivaracetam changes rapidly following intravenous injection, and it is not entirely clear how to best map occupancies to such dynamic plasma levels. Because we saw a very rapid displacement of [^11^C]UCB-J, we used peak plasma values, i.e. plasma values immediately following injection, as they align temporally to radiotracer displacement. In basement-block experiments, where the drug is normally administered some time before the block scan, the plasma drug concentration will remain relatively constant during the scan. In these cases, the plasma concentration either at the start of the scan, the end of the scan, or the mean of the two, is often used in the Emax model.^37^ For comparison, we re-ran the dose-occupancy analysis using mean plasma concentrations during each scan in place of the peak plasma values. This resulted in an *IC*_*50*_ of 0.47 μg/mL, for both the numerical solution and single-step approximation. This is nearly identical to the brivaracetam *IC*_*50*_ reported by Finnema and colleagues (0.46 μg/mL) from a [^11^C]UCB-J baseline-block experiment in humans.^38^

When performing drug development studies, it is common practice to use a range of doses to better characterise the dose-occupancy relationship. The displacement models presented here do not necessarily provide good estimates in the low occupancy ranges (∼25%, or lower). At 25% occupancy, there is also a large uncertainty in the *V*_*ND*_ estimate (supplementary figure 8), and in several cases (especially for the numerical solution) it hits the lower bound at *V*_*ND*_ = 0. Due to a strong positive correlation between *∂*^*max*^ and *V*_*ND*_ (see supplementary tables 1 and 2, and supplementary figures 11 and 12), this leads to the apparent negative bias in occupancy that we see in figure 2. Difficulties in determining low occupancies has also been reported with the Lassen plot.^39–41^ A possible solution is to fit multiple subjects simultaneously in a multilevel pharmacokinetic modelling framework, allowing the model to differentiate between displacement and normal scans.^42^ This could improve the occupancy estimates, even if normal and displacement scans were conducted in different research subjects. Such an approach could be particularly valuable if the displacement is small, e.g., when using a behavioural task to elicit neurotransmitter release rather than a pharmacological challenge.

The assumption of instantaneous occupancy (single step) that we have employed to allow the model to be solved analytically has already been shown to be a useful one for reference region quantification of displacement scans.^36^ We emphasize that although the single step solution involves splitting TACs into two segments, each segment is not fitted independently. All rate constants are constrained to be constant throughout the scan, and they are estimated by fitting the entire TAC.

The objective of introducing a numerical solution, accounting for the time course of occupancy, was to allow better quantification of occupancy for slow-acting drugs. Unexpectedly, the two approaches performed comparably across all experiments, even for the simulated slower drug (*t*_*e*_ = 30 min). In addition to the presented data, we simulated scans with a much slower drug (*t*_*e*_ = 120 min, see supplementary figure 5), where the final occupancy was reached 30 min after the last acquired data. Even in this case, we saw no advantage of the numerical solution over the single step simplification. In fact, both approaches performed poorly in this scenario.

The performance of the single step solution presented in this paper relies on using a relatively high time resolution (0.5 s frequency) in the convolution step. In our experience, this is a much shorter step size than what is usually used when solving compartment models. Consequently, the single step approach is computationally relatively heavy, and requires approximately 30 times longer run-time than the Euler Forward-based numerical solution.

Another advantage of the numerical approach is that it allows flexibility. In this study, we have used a monotonic increasing function to explain the time course of occupancy. These assumptions appear to be reasonable for the [^11^C]UCB-J pig scans with brivaracetam intervention. However, depending on factors like the drug, radiotracer, and experimental design, in some cases it might be preferable to use a different type of function to describe the occupancy. For instance, the numerical approach allows for an occupancy model where both drug uptake and washout happen during the scan. We chose the occupancy model in (equation 1) because it is a continuous and differentiable function that allows some key parameters to be estimated. In reality, we expect that the increase in occupancy in the time following a drug intervention is at first rapid, and then slows down as fewer binding sites remain available. In a recent study, Naganawa and colleagues present a similar displacement model for [^11^C]UCB-J. In that study, the rate constants defining the drug’s uptake and clearance in tissue were estimated, together with the radioligand’s rate constants.^43^ While this model more accurately reflects the underlying competition at the SV2A binding sites, the authors show that parameter identifiability becomes challenging with a model of that complexity. The approach presented in the current study is thus a pragmatic solution to derive occupancy estimates from a single displacement scan.

We are confident that the simulated TACs have a level of noise that is realistic to [^11^C]UCB-J. The level of noise could however vary for different radiotracers. In supplementary figures 6 and 7 we show histograms of *∂*^*max*^ estimates at different levels of noise. With increasing noise, the precision of the parameter estimation is reduced, but the estimates remain unbiased. With no noise added, both solutions to the model consistently return the true *∂*^*max*^ value.

In both solutions to the model, the time of the intervention (*t*_*b*_) is treated as a known parameter and is not fitted. In the presented results, the models were solved with the true *t*_*b*_ values. In reality, it might be difficult to identify the exact moment when the drug reaches its target. We therefore applied the models to some of the simulated data with wrong values for *t*_*b*_ (1 and 5 minutes before and after the true *t*_*b*_). For both solutions to the model, the *∂*^*max*^ and *V*_*ND*_ were generally unaffected by the different values for *t*_*b*_ (supplementary figure 16).

A limitation of the models is that they only consider a change in available binding sites. Pharmacological interventions may also affect perfusion, which could influence some model parameters (e.g., *K*_*1*_ for highly permeable radiotracers). If the intervention causes, for instance, an increase in perfusion, the models presented here are likely to underestimate the occupancy due to *K*_*1*_ being fixed throughout the scan. Further work is needed to develop models that can account for other changes than a reduction in *B*_*avail*_ induced by the pharmacological challenge, like some of the existing reference region based methods do.^14,19^ Also, similar to available methods for baseline-block scans, the model does not account for specific binding of the radiotracer to the target, assuming that it is only present in tracer doses. Depending on the specific activity of the radiotracer, this could lead to some bias.

Although we derived displacement versions of both the 1TCM and 2TCM (see supplementary material, section A.), we only considered a tracer that can be described by 1TC kinetics. Additional work is needed to evaluate the performance of the 2TC displacement models, as well as reference tissue implementations.

A limitation of the simulation experiments is that, for the numerical solution, the same model is used both to simulate the data and solve it. This could offer an unfair advantage to the numerical solution over the single step approximation, but our pig experiments confirm that the two approaches perform well, and they are in agreement with the Lassen plot outcome. We also limited our case to a drug that after intravenous injection shows a very immediate interaction with the target. Future studies must show if the two methods perform equally well for more slow-acting drugs. For solving the proposed displacement model, both with the numerical approach and the analytical approximation, it is necessary to pool data from several brain regions. This is standard for methods of estimating occupancies in the absence of a reference region.^5,27,39–41^

In conclusion, drug displacement PET scans constitute a promising alternative to determine occupancy, compared to baseline and follow-up studies. The kinetic models presented here enable estimation of occupancy from a single displacement scan, thereby obviating the need for two consecutive scans. This allows the number of scans required for target engagement studies to be substantially reduced, leading to lower radiation exposure and experimental costs, while also limiting the variation of biological and experimental factors. To facilitate the implementation of these models in other research centres, the MATLAB code is freely available for download at https://github.com/Gjertrud/ISI.

## 5. Acknowledgement

This project has received funding from the European Union’s Horizon 2020 research and innovation programme under the Marie Sklodowska-Curie grant agreement No 796759. GLL was supported by Rigshospitalet’s Research Council (R197-A8956). PPS was supported by the Swedish Society for Medicine and the Lundbeck Foundation (R303-2018-3263). AJ was supported by The Independent Research Fund Denmark. NRR was funded by the European Union’s Horizon 2020 research and innovation programme under the Marie Sklodowska-Curie grant agreement No 813528. HDH was supported by the Lundbeck Foundation (R293-2018-738). RTO and AAL were supported by visiting professorships from the Lundbeck Foundation (R317-2019-787 and R290-2018-756). The authors would like to thank the veterinarians and staff at the Department of Experimental Medicine, University of Copenhagen for their continued assistance with animal experiments.

## 6. Author contribution statement

MS, PPS, AAL, RTO, GMK and CS developed the theoretical framework. GLL and MS implemented the models, designed the simulations, and performed all calculations. AJ, NRR, CAM, LLD and HDH planned and conducted all PET experiments. AN analyzed all blood and plasma samples. JM led all radioligand syntheses. GLL and MS drafted the manuscript, and all authors provided critical feedback and helped shape the final version.

## 8. Supplementary material

Supplementary material can be found in a separate file.

