## Supplemental material for "Kinetic models for PET displacement studies"

### A. Displacement model for 2TCM kinetics

#### A.1 Operational equations for the displacement model based on 2TCM kinetics

In the 2TCM, the rate of exchange between compartments is determined by the constants *K_1_*, *k_2_*, *k_3_* and *k_4_* (Figure 1). The rate constant *k_3_* is linearly dependent on the concentration of available targets ($B_{avail}$), $k_{3}=f_{ND}k_{on}B_{avail}$.^1,2^ We assume that a reduction of available targets will have a negligible impact on both the association rate constant $k_{on}$ and the fraction of free radioligand in the non-displaceable compartment, $f_{ND}$. It follows that a time dependent reduction of available targets, i.e., $(1-\partial\left( t \right))B_{avail}$ will affect *k_3_* equally, i.e., $f_{ND}k_{on}\cdot\left( 1-\partial\left( t \right) \right)B_{avail}={(1-\delta\left( t \right))k}_{3}$. With this, the 2TCM can be modified to accommodate an increase in occupancy, starting at some time $t_{b}$ after radioligand injection. A schematic diagram of the model is shown in Figure 1, with the following differential equations:

$\left\{ \begin{aligned} \frac{dC_{ND}(t)}{dt}=K_{1}C_{p}\left( t \right)-\left( k_{2}+\left( 1-\partial\left( t \right) \right)k_{3} \right)C_{ND}\left( t \right)+k_{4}C_{s}(t) \\ \frac{dC_{S}(t)}{dt}={\left( 1-\partial\left( t \right) \right)k}_{3}C_{ND}\left( t \right)-k_{4}C_{s}\left( t \right) \end{aligned} \right.$,	(s1)

where $C_{p}\left( t \right)$is the metabolite corrected arterial plasma input function, $C_{s}$ and $C_{ND}$ are radioligand concentrations in the respective compartments, and

$\partial\left( t \right)=\left\{ \begin{aligned} 0, t<t_{b} \\ \partial^{max}\left( 1+\frac{t_{e}-t}{t_{e}-t_{m}} \right)\left( \frac{t-t_{b}}{t_{e}-t_{b}} \right)^{\frac{t_{e}-t_{b}}{t_{e}-t_{m}}}, t_{b}\leq t<t_{e} \\ \partial^{max}, t\geq t_{e} \end{aligned} \right.$	.	(s2)

#### A.2 Single step approximation for the 2TCM

Let $C_{T}^{0}\left( t \right)$and $C_{T}^{1}\left( t \right)$ denote tissue concentrations before and after administration of the competing drug, respectively. The equations describing $C_{ND}^{0}\left( t \right)$ and $C_{S}^{0}\left( t \right)$ are the standard differential equations for the 2TCM. By introducing a new time variable, $\tau=t-t_{s},$ the differential equations for $t\geq t_{s}$ become

$\left\{ \begin{aligned} \frac{dC_{ND}^{1}(\tau)}{d\tau}=K_{1}C_{p}\left( \tau+t_{s} \right)-\left( k_{2}+\left( 1-\partial^{max} \right)k_{3} \right)C_{ND}^{1}\left( \tau\right)+k_{4}C_{S}^{1}\left( \tau\right) \\ \frac{dC_{S}^{1}\left( \tau\right)}{d\tau}={\left( 1-\partial^{max} \right)k}_{3}C_{ND}^{1}\left( \tau\right)-k_{4}C_{S}^{1}\left( \tau\right) \\ C_{ND}^{1}\left( t_{s} \right)= C_{ND}^{0}\left( t_{s} \right), C_{S}^{1}\left( t_{s} \right)= C_{S}^{0}\left( t_{s} \right) \end{aligned} \right.$, $\tau\geq0$	(s3)

Following the nomenclature provided in Gunn et al., ^3^ the analytical solution for total activity in tissue, $C_{T}\left( t \right)=C_{ND}\left( t \right)+C_{S}\left( t \right)$, becomes

$$C_{T}\left( t \right)=$$

$$\left\{ \begin{matrix} C_{p}\left( t \right)\otimes\left( \phi_{1}e^{-\theta_{1}t}+\phi_{2}e^{-\theta_{2}t} \right), t\leq t_{s} \\ \begin{matrix} C_{p}\left( t \right)\otimes\left( \phi_{1}e^{-\theta_{1}\left( t-t_{b} \right)}+\phi_{2}e^{-\theta_{2}\left( t-t_{b} \right)} \right)+C_{ND}^{0}\left( t_{b} \right)\frac{\left( \phi_{1}e^{-\theta_{1}\left( t-t_{b} \right)}+\phi_{2}e^{-\theta_{2}\left( t-t_{b} \right)} \right)}{K_{1}} \\ +C_{S}^{0}\left( t_{b} \right)\left( \frac{-\theta_{2}e^{\theta_{1}\left( t-t_{b} \right)}+\theta_{1}e^{\theta_{2}\left( t-t_{b} \right)}}{\Delta} \right) \end{matrix}, t>t_{s} \end{matrix} \right.$$

(s4)

Expressions for $\phi_{i}$, $\theta_{i}$, $\Delta$, $C_{ND}^{0}\left( t_{b} \right)$ and $C_{S}^{0}\left( t_{b} \right)$ are,

$C_{ND}^{0}\left( t_{b} \right)= \left( H_{ND}\bigotimes C_{p} \right)\left( t_{b} \right)$

$C_{S}^{0}\left( t_{b} \right)= \left( H_{S}\bigotimes C_{p} \right)\left( t_{b} \right)$

$$\left\{ \begin{matrix} \phi_{1}=\frac{K_{1}\left( \theta_{1}-{\left( 1-\partial\right)k}_{3}{-k}_{4} \right)}{\Delta} \\ \phi_{2}=\frac{K_{1}\left( \theta_{2}-{\left( 1-\partial\right)k}_{3}{-k}_{4} \right)}{-\Delta} \\ \theta_{1}\boldsymbol{=}\frac{k_{2}+{\left( 1-\partial\right)k}_{3}+k_{4}+\Delta}{2} \\ \theta_{2}\boldsymbol{=}\frac{k_{2}+{\left( 1-\partial\right)k}_{3}+k_{4}-\Delta}{2} \\ \Delta\boldsymbol{=}\sqrt{\left( k_{2}+\left( 1-\partial\right)k_{3}+k_{4} \right)^{2}-4k_{2}k_{4}} \\ H_{ND}\left( t \right)=\frac{K_{1}}{\Delta}\left( \left( \theta_{1}-k_{4} \right)e^{-\theta_{1}t}-\left( \theta_{2}-k_{4} \right)e^{-\theta_{2}t} \right) \\ H_{S}\left( t \right)= \frac{K_{1}k_{3}}{\Delta}\left( -e^{-\theta_{1}t}+e^{-\theta_{2}t} \right) \end{matrix} \right.$$

where,

$\partial=\left\{ \begin{aligned} 0, t<t_{b} \\ \partial^{max}, t\geq t_{b} \end{aligned} \right.$ ,

#### A.3 Numerical solution for the 2TCM

The Euler Forward Numerical solution for the 2TCM (i.e., equation s1) becomes

$\left\{ \begin{aligned} C_{ND}\left( t_{n} \right)=hK_{1}C_{p}\left( t_{n-1} \right)+\left( 1-h\left( k_{2}+{\left( 1-\partial\left( t_{n-1} \right) \right)k}_{3} \right) \right)C_{ND}\left( t_{n-1} \right)+hk_{4}C_{s}\left( t_{n-1} \right) \\ C_{s}\left( t_{n} \right)=\left( 1-hk_{4} \right)C_{S}\left( t_{n-1} \right)+h\left( 1-\partial\left( t_{n-1} \right) \right)k_{3}C_{ND}\left( t_{n-1} \right) \end{aligned} \right.$.

(s5)

With the occupancy function $\partial(t)$ according to equation s2.

### B. Rationale for simultaneously fitting several ROIs

As mentioned in the main text (section 2.1.4), the model parameters $k_{2}$, $\partial(t)$, and ${BP}_{ND}$ appear only as a ratio in the proposed model, and as a result, these parameters cannot be uniquely identified when the model is applied to a single TAC. This is illustrated in Supplementary figure 1, which shows surface plots of the model objective function for varying values for *V_ND_* and occupancy, when the model is applied to each region separately. For all regions, there is a line of combinations of *V_ND_* and occupancy that yield similarly low values of the objective function. Supplementary figure 2 shows the profile of the objective function through those lines. The profile does not have a clear global minimum for any of the regions. However, when all the profiles are combined, we get a clear minimum at *V_ND_* = 4, which is the true value for *V_ND_* in the simulations.

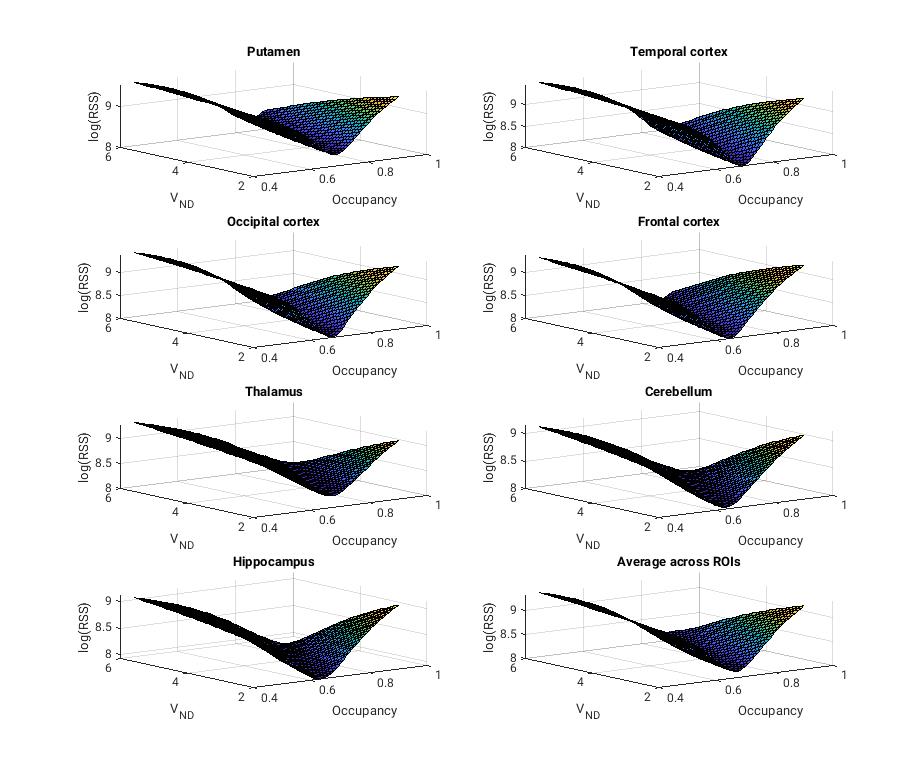

Supplementary figure 1: Objective function of model applied to a single TAC, plotted against V_ND_ and ∂^max^ (occupancy). ROIs are (left to right, top to bottom) putamen, temporal cortex, occipital cortex, frontal cortex, thalamus, cerebellum, and hippocampus. The final plot (bottom right) is the average across all ROIs.

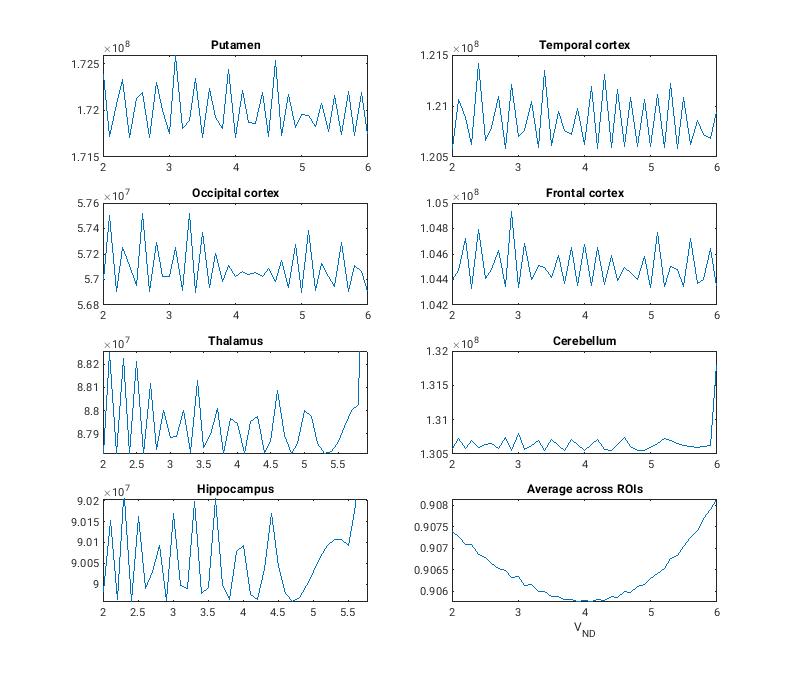

Supplementary figure 2: Line profiles through the valleys of the objective functions in Supplementary figure 1. ROIs are (left to right, top to bottom) putamen, temporal cortex, occipital cortex, frontal cortex, thalamus, cerebellum, and hippocampus. The final plot (bottom right) is the average across all ROIs.

### C. Implementation of models

All analyses were performed in Matlab (version 9.10). For both solutions to the model the fitting was run with a nested approach, where the outer layer function fitted the global parameters (*∂^max^*, and *V_ND_* for both solutions, *t_e_* for the numerical solution, and *t_s_* for the single step approximation). For each iteration of the outer function, the ROI-specific parameters (*V_S_*, *K_1_* and *v_B_*) were fitted for each ROI separately. In both layers of the algorithm, lsqnonlin was used to fit the model to the data. *∂^max^* and *v_B_* were constrained to be between 0 and 1. *t_e_* and *t_s_* were constrained to be higher than (after) *t_b_*. All other parameters were constrained to be positive.

Matlab code for both models is freely available and can be found at <https://github.com/Gjertrud/ISI>.

### D. Blood and plasma analyses

The blood samples were centrifuged (2246xg for 7 min at 4°C), and the extracted plasma was filtered through a 0.45 µm syringe filter (Whatman GD/X 13 mm, Cytiva) and subsequently diluted 1:1 with 20 mM Phosphate buffer and 5 mM sodium-1-decanesulfonate pH 7.2 with 2% isopropanol. Samples were analysed in a fully automated column-switching HPLC system (UltiMate 3000, Thermo Fisher Scientific) connected to a radio-HPLC detector (PosiRam Model 4, LabLogic Systems).^4^ The HPLC system was equipped with a small extraction column (Shimpack MAYI-ODS 30x4.6 mm, Shimadzu Corporation) combined with an analytical column (Onyx Monolithic C18 50x4.6 mm, Phenomenex). The extraction mobile phase consisted of 100% of phosphate buffer (composition mentioned above), while the elution mobile phase consisted of 59% of 100 mM phosphate buffer and 2 mM sodium 1-decanesulfonate pH 2.6 and 41% methanol. Samples were injected in a volume of 4 mL, and the analysis was run at a flow of 5 mL/min at 25 ºC. Total runtime for each sample was 8.55 min with a 4 min extraction step, 4 min elution step and 0.55 min of equilibration. Four eluate fractions were collected in 2 min intervals using a fraction collector (Foxy Jr FC144; Teledyne) and radioactivity subsequently measured using a gamma counter (Wizard 2480, Perkin Elmer). The parent tracer fraction was calculated as follow: % parent fraction = (radioactivity of parent eluate/total amount of collected radioactivity) x 100%.

### E. Supplementary results

#### E.1 Example of simulated data

The following two figures show examples of simulated TACs. In Supplementary figure 3, four different temporal cortex TACs from the same simulation experiment (*t_e_* = 30 min, ∂^max^ = 50%, $\alpha$ = 5) are shown. Supplementary figure 4 shows five simulated temporal cortex TACs with increasing noise for a simulated displacement scan with *t_e_* = 5 and ∂^max^ = 75%.

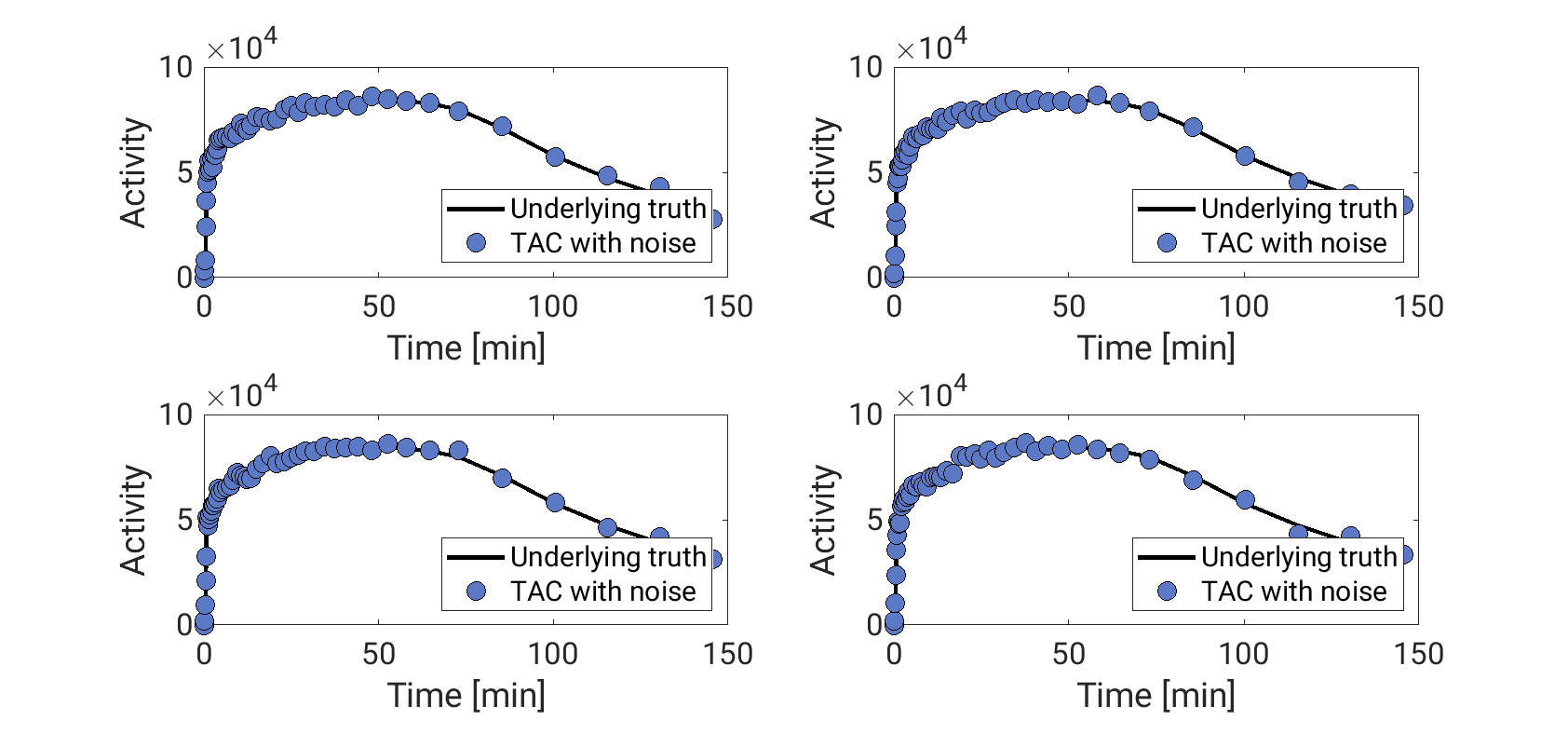

Supplementary figure 3: examples of four different simulated temporal cortex TACs for a displacement scan where 50% occupancy is reached 30 min after the intervention. The noise level ($\alpha$) was 5.

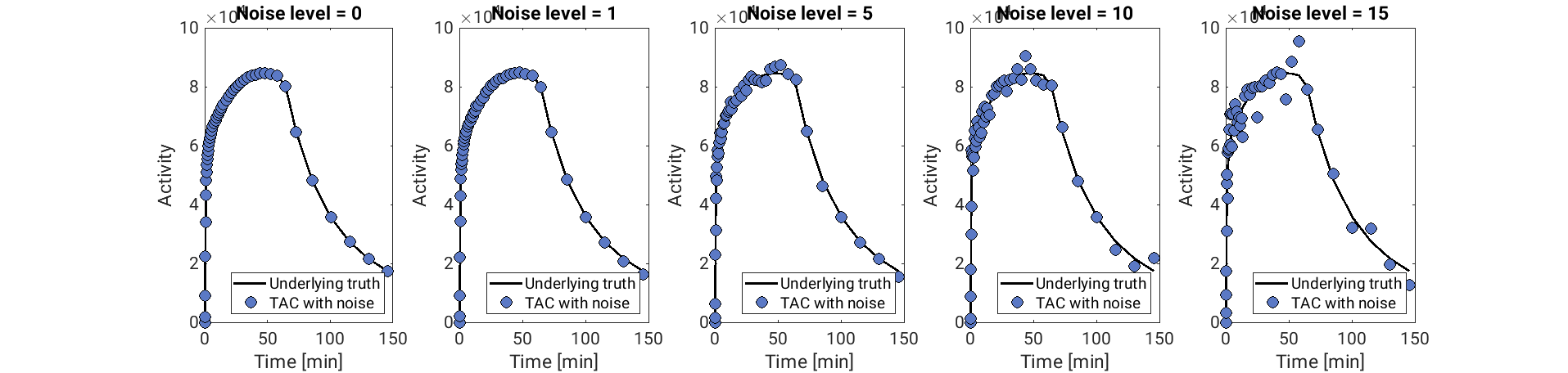

Supplementary figure 4: examples of simulated temporal cortex TACs at five different noise levels ( α = 0, 1, 5, 10 & 15) for a displacement scan where 75% occupancy was reached after 5 min.

#### E.2 Results for simulated drug with t_e_ = 120

In the main text we show parameter estimates for displacement scans where full occupancy (∂^max^) was reached 5 or 30 minutes after the intervention. In addition, we simulated displacement TACs where full occupancy was reached after 120 minutes, i.e., 30 minutes after the end of the PET scan. The results from this experiment are illustrated in Supplementary figure 5, with ∂^max^ estimates on the top row and *V_ND_* estimates on the bottom row. For all occupancies, and both model solutions, the fitting generally failed. The single-step solution displays a tendency to underestimate ∂^max^ (by approximately half). The numerical solution resulted in very wide distribution of ∂^max^ estimates, frequently landing on the upper limit of 100%.

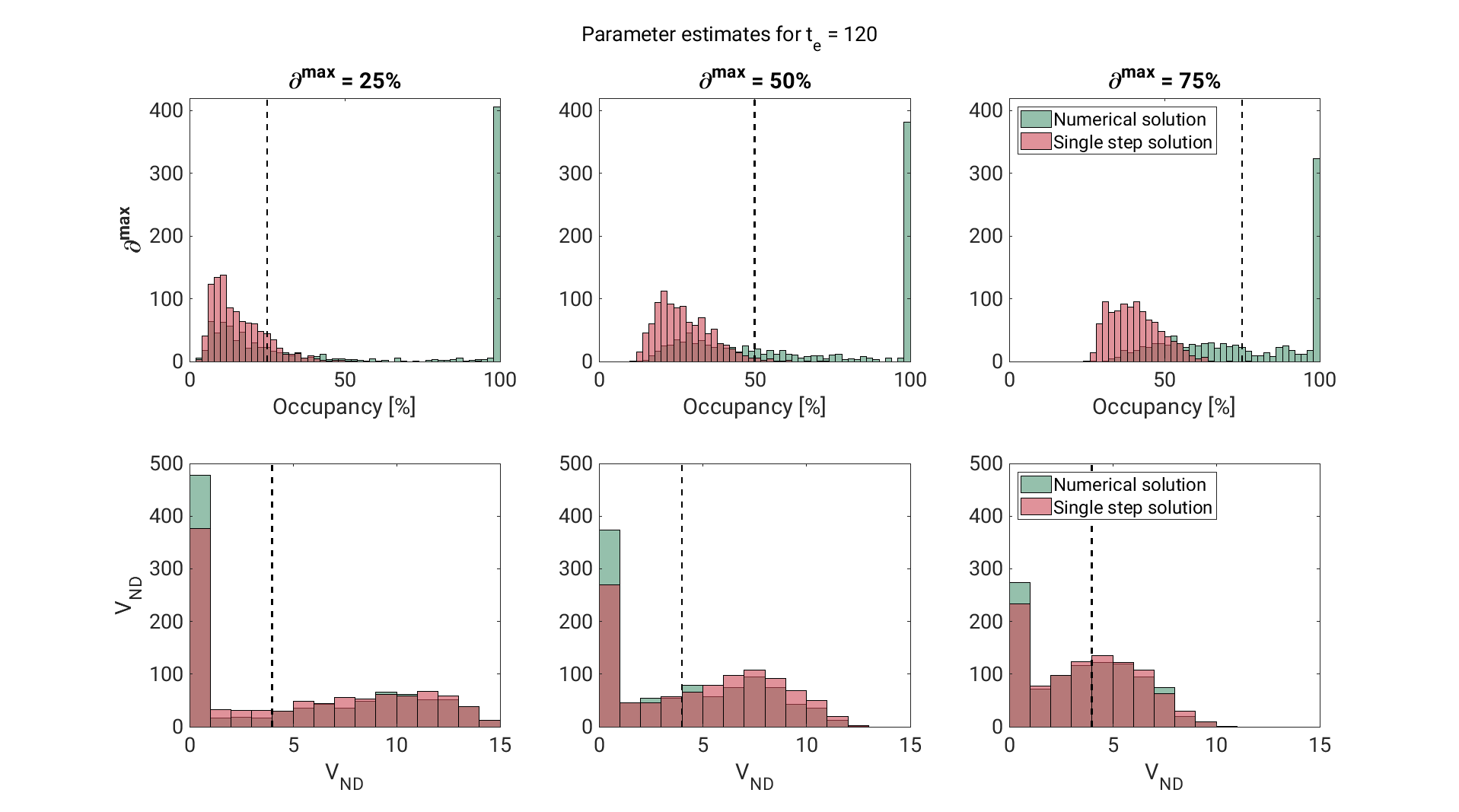

Supplementary figure 5: histograms of parameter estimates (occupancy (∆), and V_ND_) from a simulation experiment where full occupancy is reached after 120 minutes. The dashed lines represent the true parameter values.

#### E.3 Effect of noise on parameter estimates

The following figures show histograms of ∂^max^ estimates from a simulation experiment with ∂^max^ = 75% and *t_e_* = 5 min, with increasing noise added to the TACs (α = 0, 1, 5, 10 and 15). Supplementary figure 6 shows histograms for all five noise levels, and Supplementary figure 7 shows histograms for only the three highest noise levels (α = 5, 10, and 15). At α = 0 (no noise), both solutions consistently returned the true occupancy value. While the precision of the estimates deteriorated with increasing noise, the noise does not seem to induce bias.

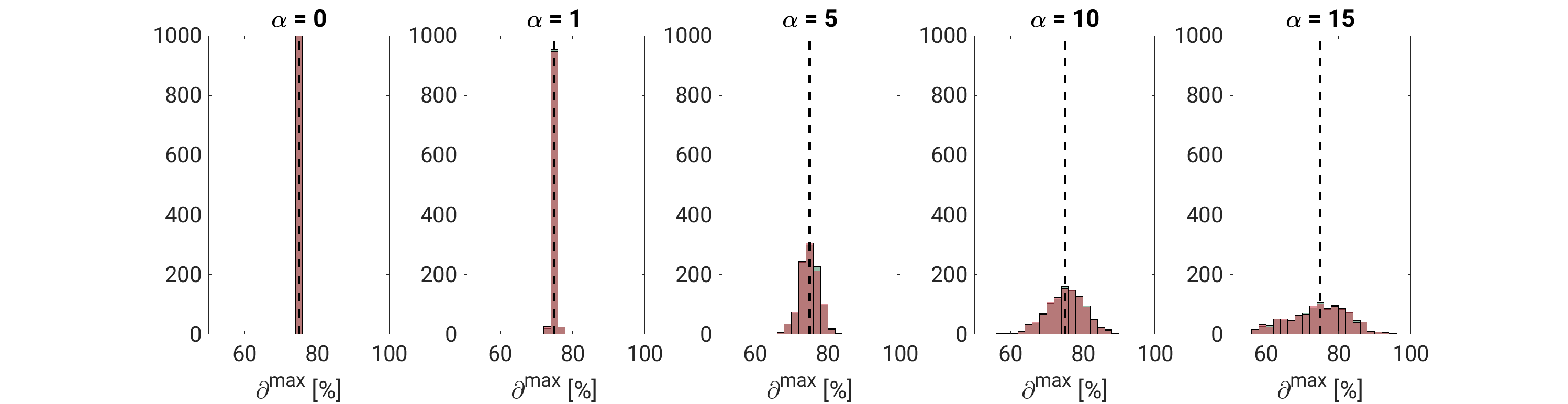

Supplementary figure 6: Histograms of occupancy estimates for increasing noise (α = 0, 1, 5, 10 & 15). In all histograms, occupancy = 75% and t_e_ = 5 min.

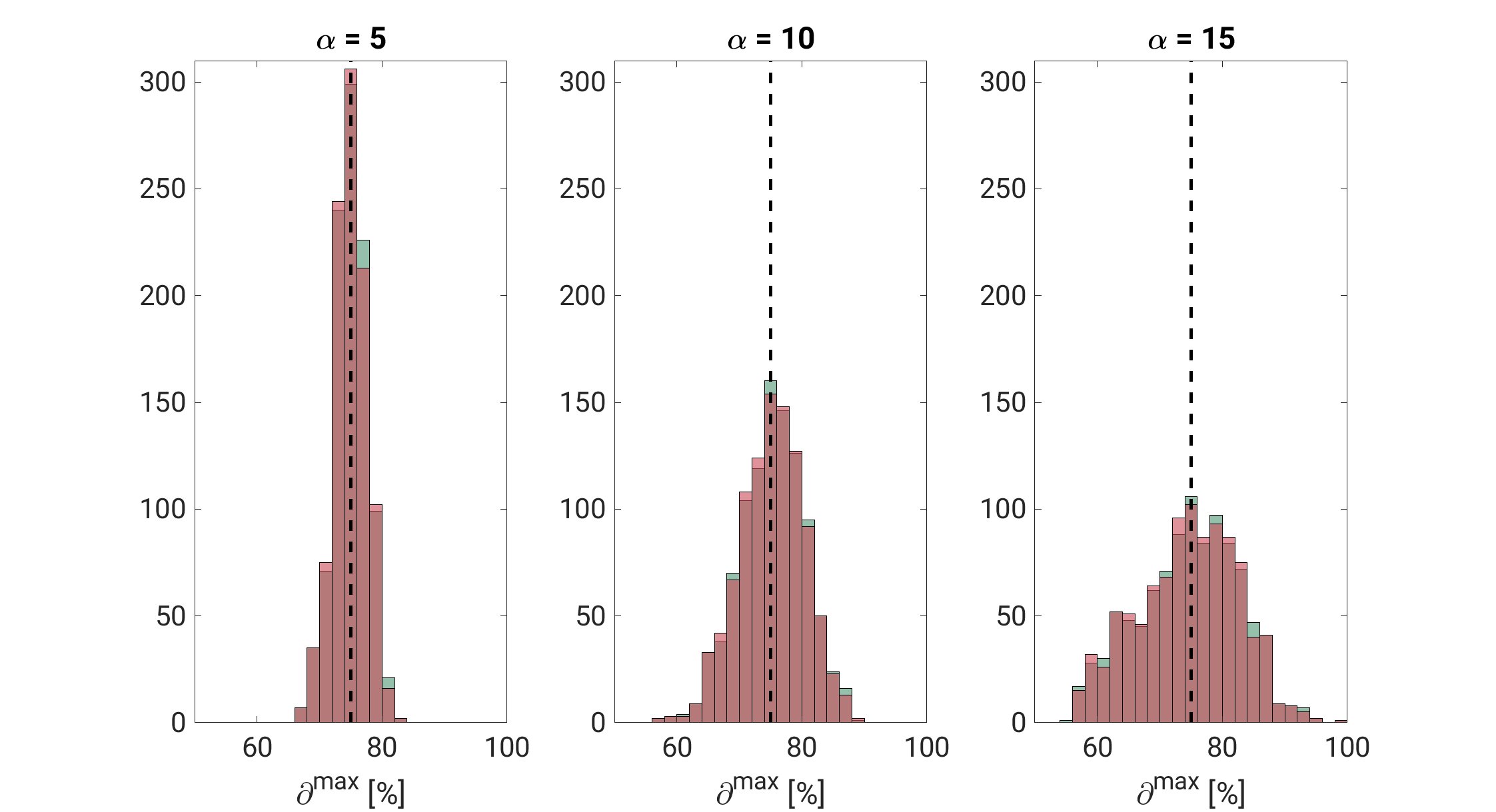

Supplementary figure 7: Histograms of occupancy estimates for increasing noise (α = 5, 10 & 15). In all histograms, occupancy = 75% and t_e_ = 5 min.

#### E.4 Estimation of V_ND_ and V_T_

The following figures show histograms of *V_ND_* (Supplementary figure 8) and temporal cortex *V_S_* (Supplementary figure 9), for ∂^max^ = 25%, 50% and 75% and *t_e_* = 5 and 30 min, estimated with both solutions to the model.

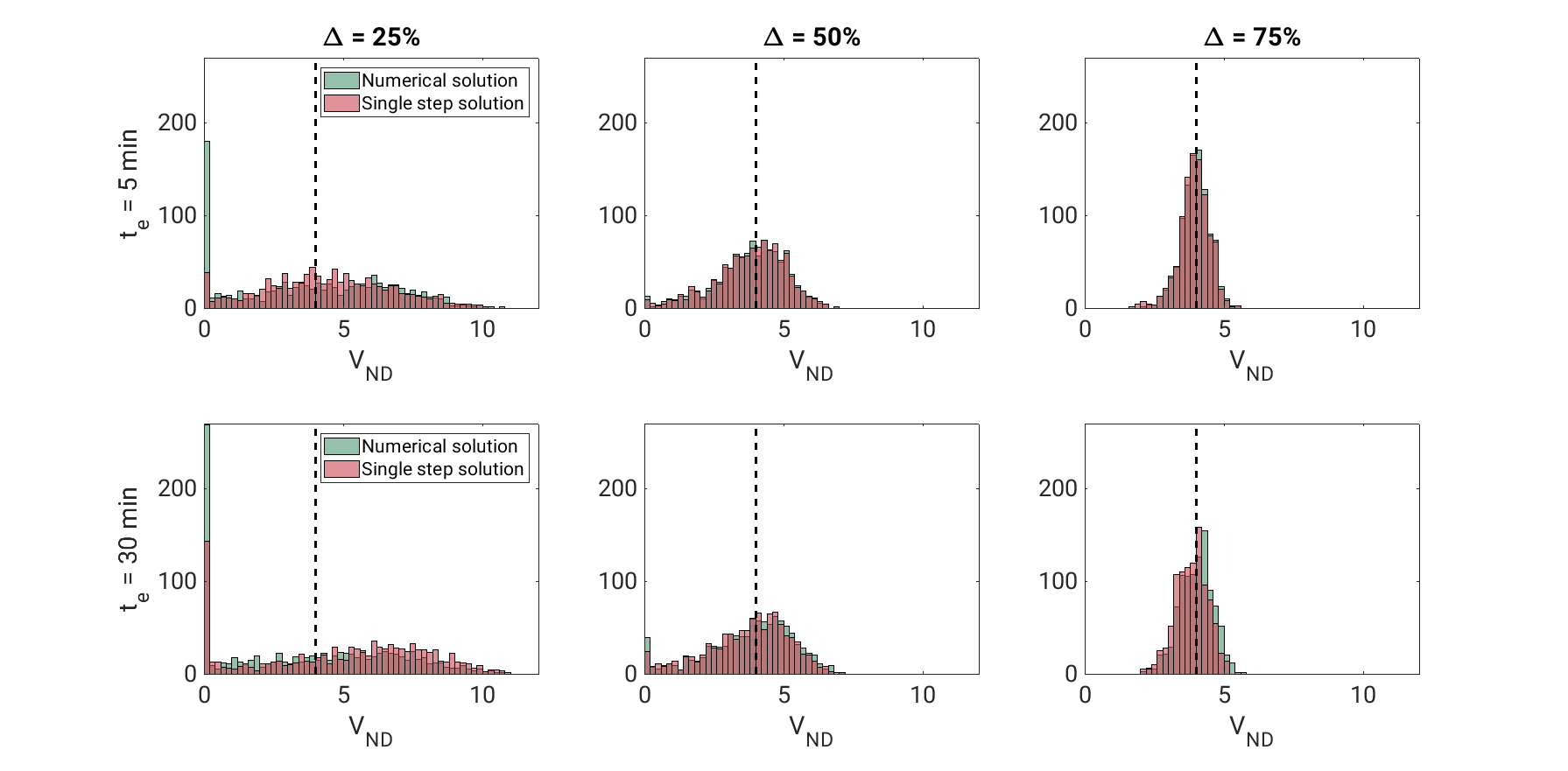

Supplementary figure 8: Histograms of V_ND_ estimates from both the numerical solution (green) and single step solution (red), for ∂^max^ = 25%, 50% and 75%, and t_e_ = 5 and 30 min.

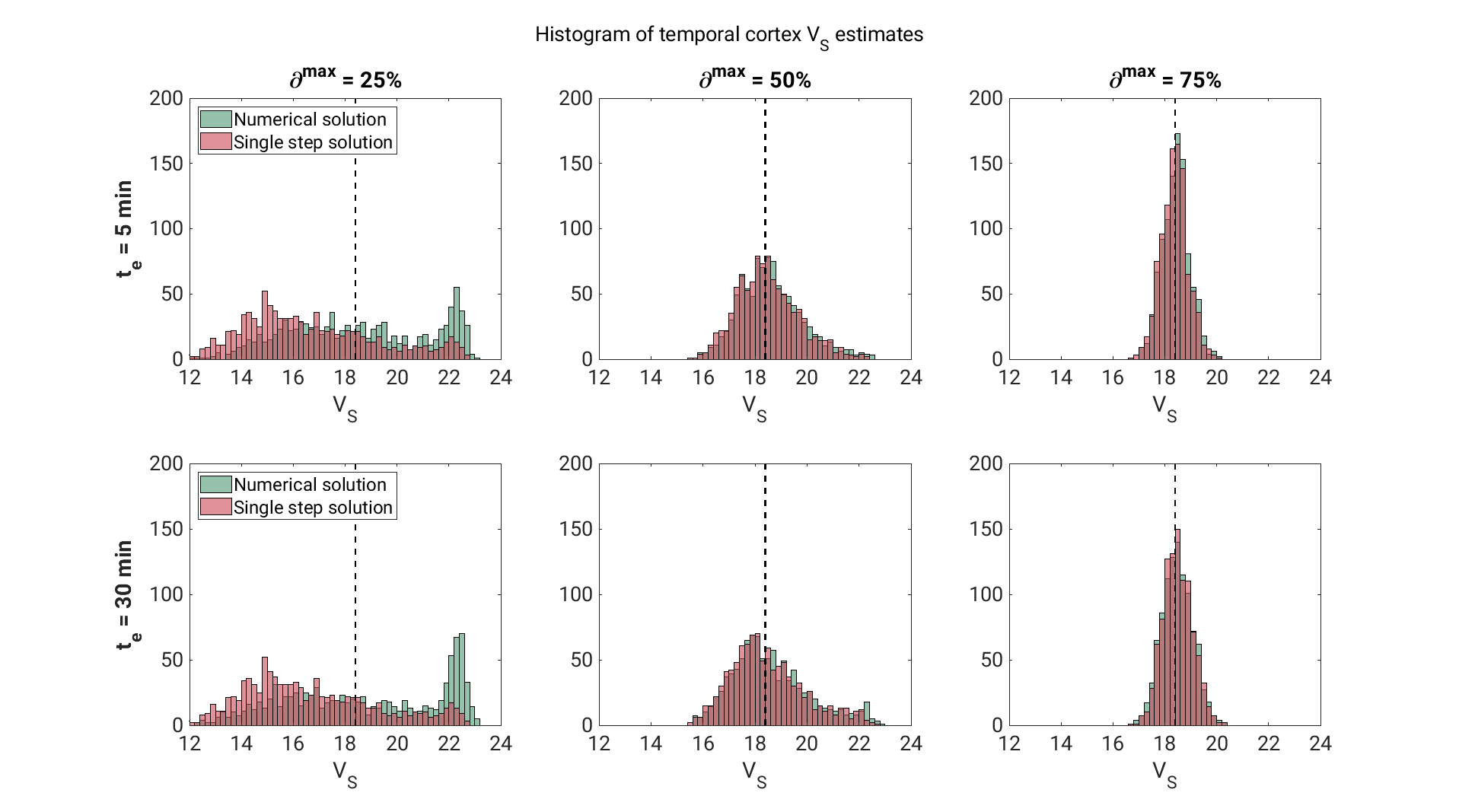

Supplementary figure 9: Histograms of temporal cortex V_S_ estimates from both the numerical solution (green) and single step solution (red), for ∂^max^ = 25%, 50% and 75%, and t_e_ = 5 and 30 min.

#### E.5 Correlation between parameter estimates

Supplementary tables 1 and 2 shows the Pearson correlation coefficients and corresponding p-values for all combinations of fitted parameters, for the numerical and single step solutions, respectively. Data from the simulations with ∂^max^ = 75% and t_e_ = 5 min was used. For ROI-specific parameters (*V_S_*, *K_1_* and *v_B_*) the temporal cortex estimates were used. Results were largely comparable between the two solutions. For the numerical solution only, scatter plots of all parameter estimates plotted against each other are presented in Supplementary figure 10 (for global parameters only) and Supplementary figure 11 (for all parameters). Again, in Supplementary figure 11, temporal cortex estimates were used for the ROI-specific parameters.

|  | *∂^max^* | *V_ND_* | *t_e_* | *V_S_* | *K_1_* | *v_B_* |
| --- | --- | --- | --- | --- | --- | --- |
| *∂^max^* |  | R = 0.98  P < 1e-100 | R = 0.047  P = 0.13 | R = -0.76  P < 1e-100 | R = -0.15  P = 1.03e-6 | R = 0.011  P = 0.74 |
| *V_ND_* | R = 0.98  P < 1e-100 |  | R = -0.011  P = 0.74 | R = -0.80  P < 1e-100 | R = -0.12  P = 9.8e-5 | R = 0.013  P = 0.68 |
| *t_e_* | R = 0.047  P = 0.13 | R = -0.011  P = 0.74 |  | R = -0.13  P = 2.2e-5 | R = 0.062  P = 0.051 | R = -0.051  P = 0.10 |
| *V_S_* | R = -0.76  P < 1e-100 | R = -0.80  P < 1e-100 | R = -0.13  P = 2.2e-5 |  | R = 0.02  P = 0.47 | R = 0.31  P < e-10 |
| *K_1_* | R = -0.15  P = 1.03e-6 | R = -0.12  P = 9.8e-5 | R = 0.062  P = 0.051 | R = 0.02  P = 0.47 |  | R = 0.57  P < 1e-10 |
| *v_B_* | R = 0.011  P = 0.74 | R = 0.013  P = 0.68 | R = -0.051  P = 0.10 | R = 0.31  P < e-10 | R = 0.57  P < 1e-10 |  |

Supplementary table 1: correlation coefficients (R) and p-values (P) for the correlations for all parameters estimated with the numerical solution. Data is taken from the simulation experiment where 75% occupancy was reached after 5 min. For ROI-specific parameters (V_S_, K_1_ and v_B_), values for temporal cortex are used. Significant correlations are marked by green backgrounds.

|  | *∂^max^* | *V_ND_* | *t_s_* | *V_S_* | *K_1_* | *v_B_* |
| --- | --- | --- | --- | --- | --- | --- |
| *∂^max^* |  | R = 0.98  P < 1e-100 | R = 0.024  P = 0.44 | R = -0.75  P < 1e-100 | R = -0.16  P = 5.8e-7 | R = 0.012  P = 0.74 |
| *V_ND_* | R = 0.98  P < 1e-100 |  | R = -0.029  P = 0.36 | R = -0.79  P < 1e-100 | R = -0.13  P = 6.3e-5 | R = 0.014  P = 0.67 |
| *t_s_* | R = 0.024  P = 0.44 | R = -0.029  P = 0.36 |  | R = -0.13  P = 2.7e-5 | R = 0.073  P = 0.020 | R = -0.050  P = 0.12 |
| *V_S_* | R = -0.75  P < 1e-100 | R = -0.79  P < 1e-100 | R = -0.13  P = 2.7e-5 |  | R = 0.020  P = 0.52 | R = 0.32  P < 1e-10 |
| *K_1_* | R = -0.16  P = 5.8e-7 | R = -0.13  P = 6.3e-5 | R = 0.073  P = 0.020 | R = 0.020  P = 0.52 |  | R = 0.57  P < 1e-10 |
| *v_B_* | R = 0.012  P = 0.74 | R = 0.014  P = 0.67 | R = -0.050  P = 0.12 | R = 0.32  P < 1e-10 | R = 0.57  P < 1e-10 |  |

Supplementary figure 10: correlation coefficients (R) and p-values for the correlations (P) for all parameters estimated with the single step solution. Data is taken from the simulation experiment where 75% occupancy was reached after 5 min. For ROI-specific parameters (V_S_, K_1_ and v_B_), values for temporal cortex are used. Significant correlations are marked by green backgrounds.

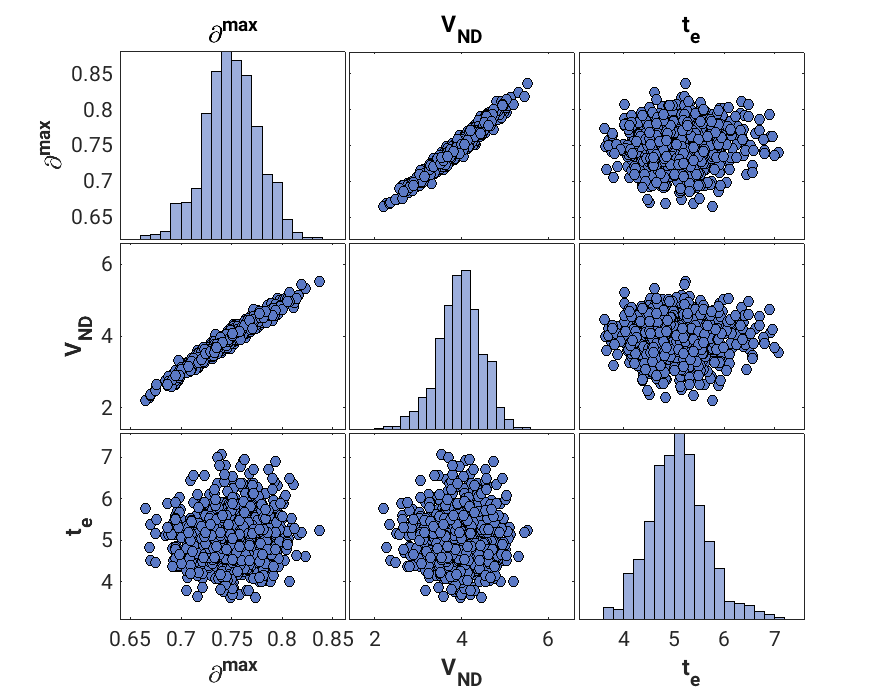

Supplementary figure 11: Scatter plots of all fitted global parameters plotted against each other. Parameters are estimated from the simulated data where ∂^max^ = 75%, and t_e_ = 5 min, using the numerical solution. On the diagonal, subplots display histograms of parameter estimates for each of the global parameters.

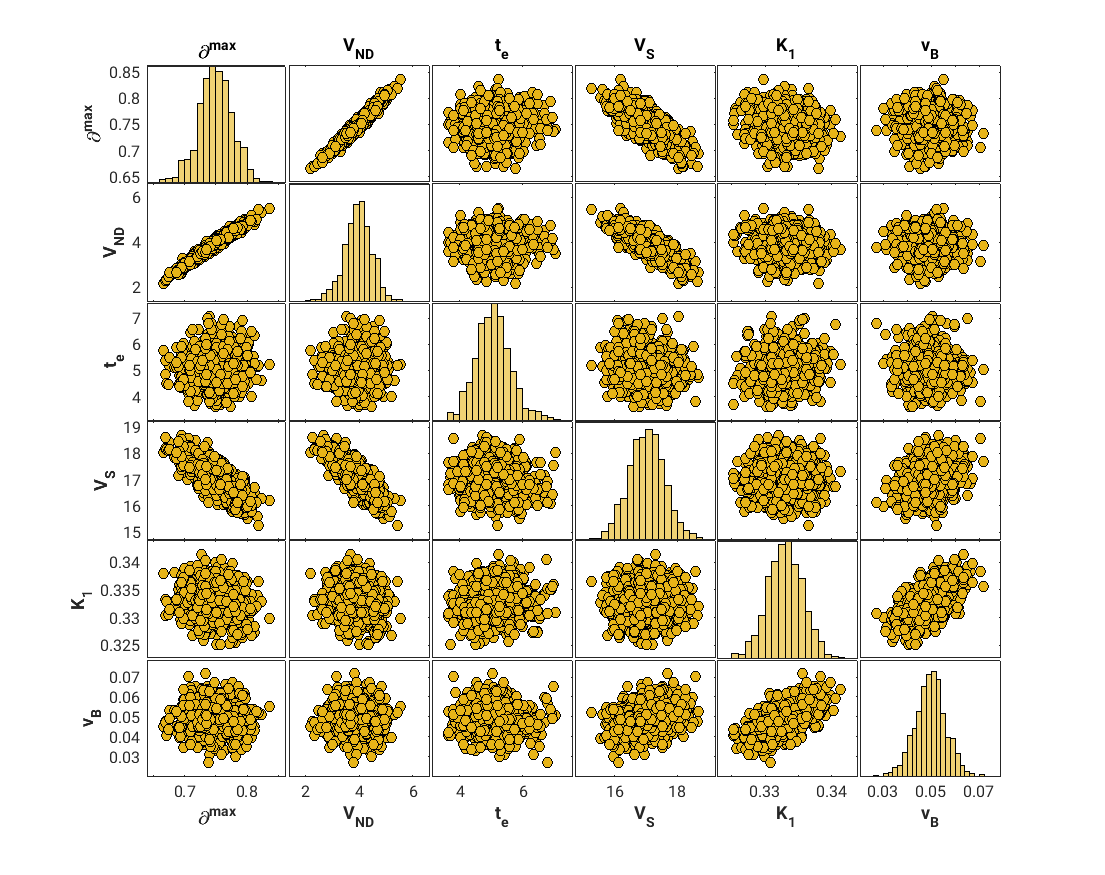

Supplementary figure 12: Scatter plots of all fitted parameters plotted against each other. Parameters are estimated from the simulated data where ∂^max^ = 75%, and t_e_ = 5 min, using the numerical solution. On the diagonal, subplots display histograms of parameter estimates for each of the parameters. For the ROI-specific parameters (V_S_, K_1_ and v_B_) estimates for temporal cortex were used.

#### E.6 Residuals of model fits to pig data

In the main text (Figure 3), we show the fits of both model solutions to temporal cortex TACs from two different pig scans with brivaracetam intervention at 60 min. Supplementary figures 13 and 14 show the model fit (numerical solution) to all included TACs in the same scans. These figures also show the residuals. In supplementary figure 15, normalized residuals for all six pig scans are shown, for putamen, temporal cortex, occipital cortex and hippocampus.

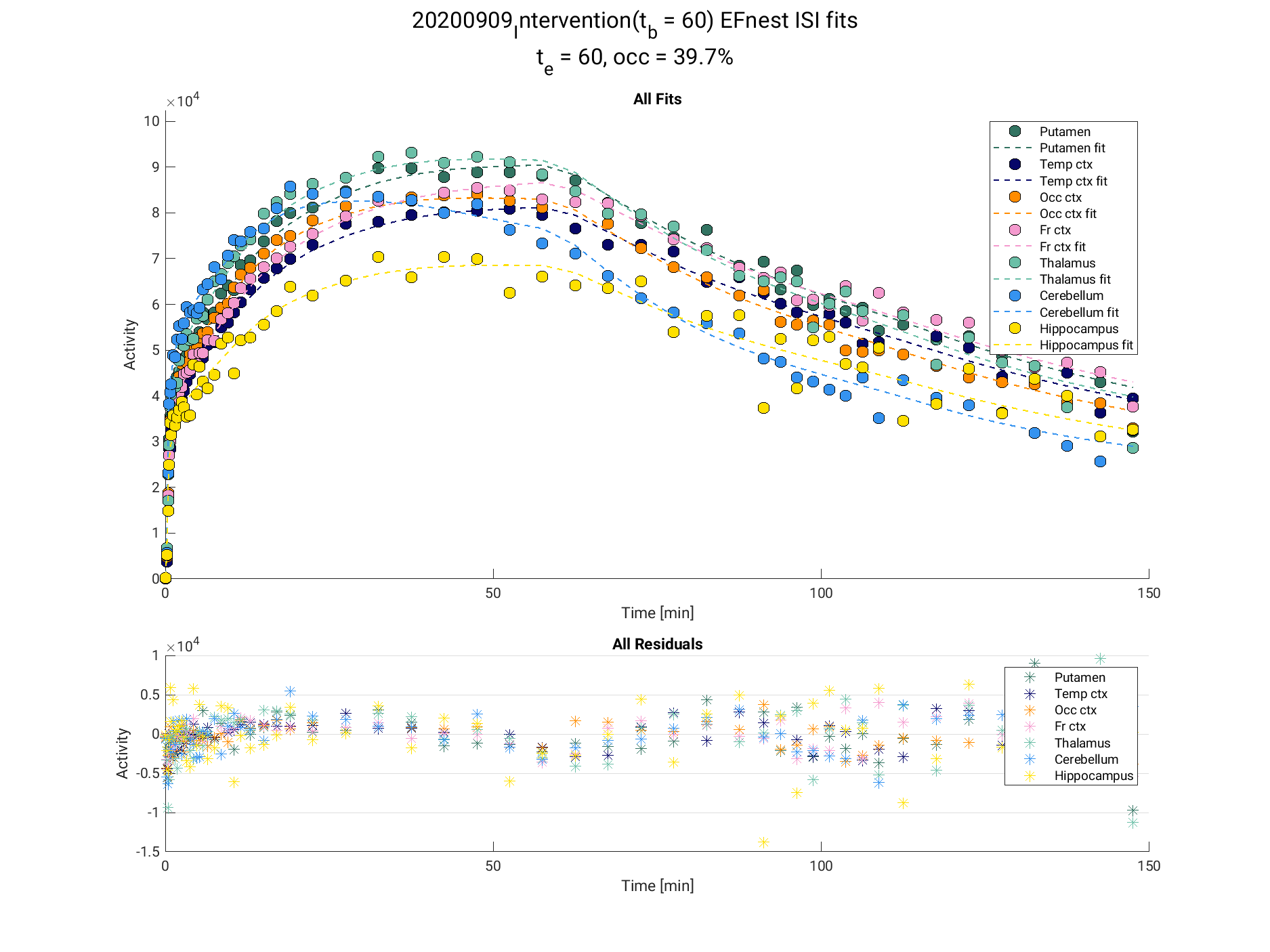

Supplementary figure 13: displacement model fits (numerical solution) to all included TACs from the pig displacement scan with the lowest brivaracetam dose (0.1 mg/kg), including residuals.

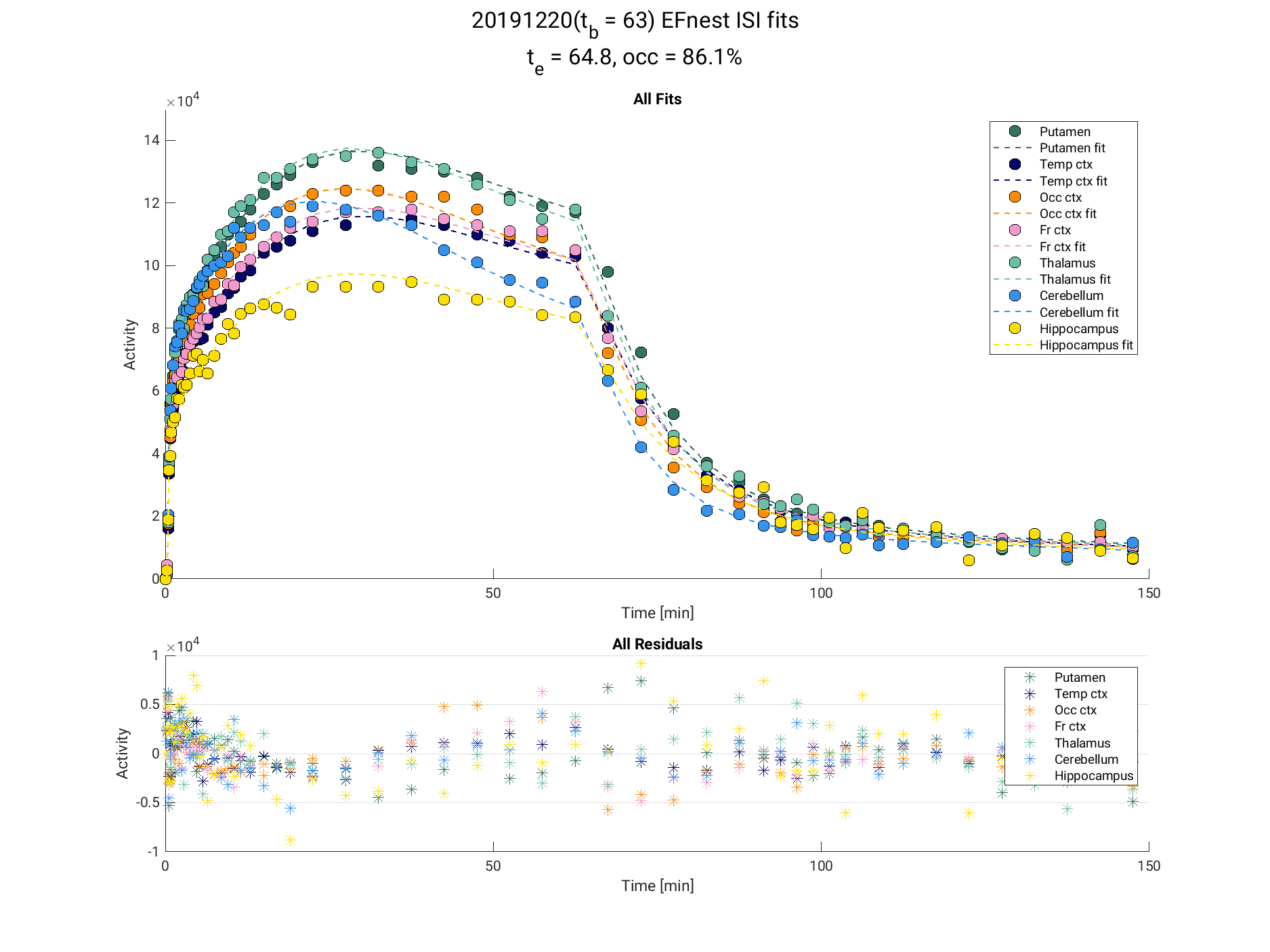

Supplementary figure 14: displacement model fits (numerical solution) to all included TACs from the pig displacement scan with the highest brivaracetam dose (5 mg/kg), including residuals.

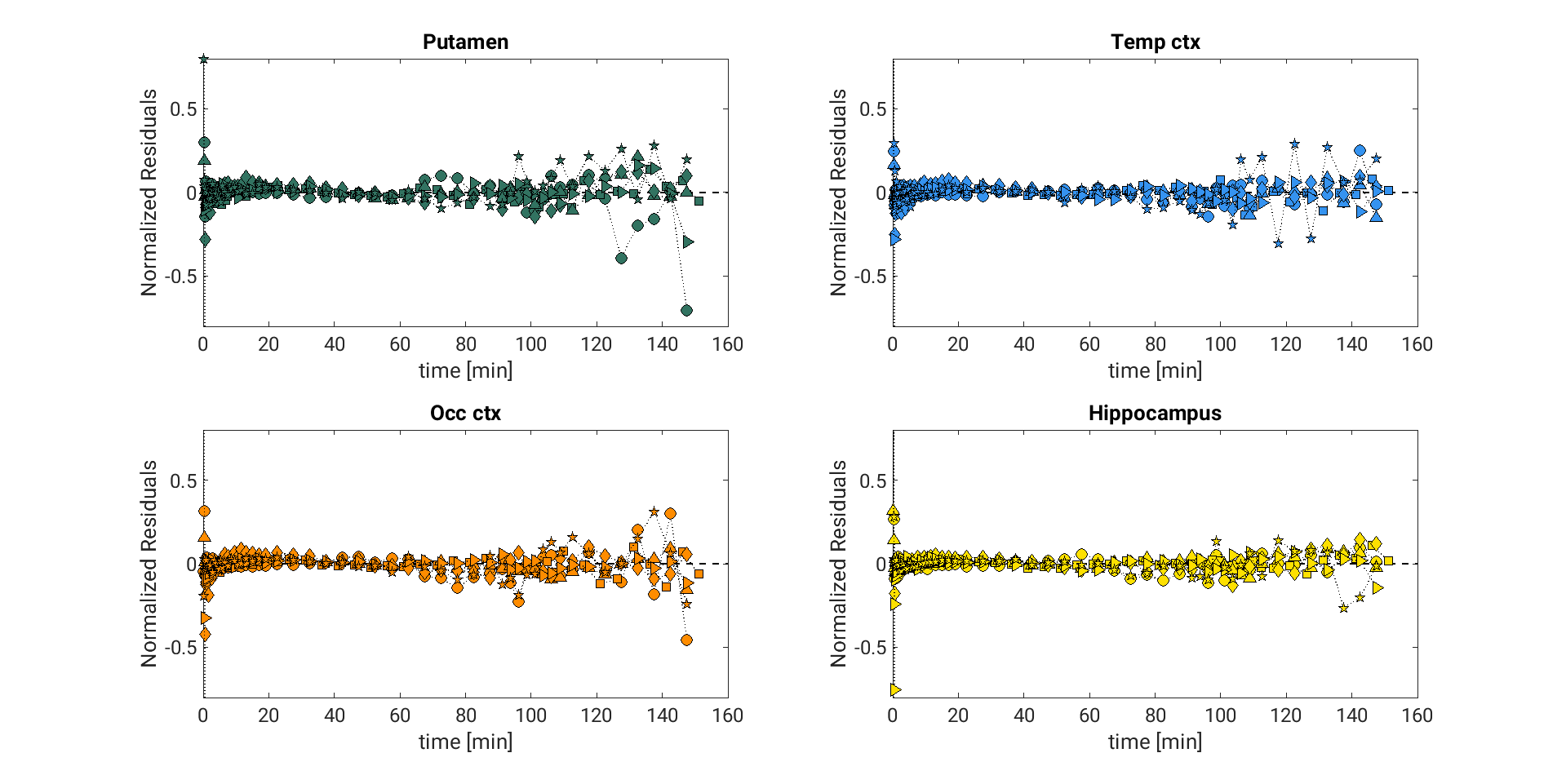

Supplementary figure 15: Normalized residuals from the model fits (numerical solution) for all six pig scans, for putamen, temporal cortex, occipital cortex and hippocampus. Each pig has a unique marker, which is consistent across the subplots. Residuals were normalized to the ROI C_T_ frame-by-frame.

#### E.7 Effect of t_b_ on parameter estimates

In the presented displacement model, t_b_ (the time of the intervention) is treated as a known parameter. In real life, the drug cannot enter the circulation instantaneously, but is typically injected over seconds-minutes. Also, there will be some delay between administration of drug, and drug reaching its brain target. Thus, choosing t_b_ is not trivial.

To investigate the effect of a badly defined t_b_, we fitted the model to the simulated data where ∂^max^ = 75% and *t_e_* = 5 min using different values for *t_b_*. The data was simulated using *t_b_* = 60 min. We solved the data with both the numerical and single step approaches, and *t_b_* defined to be 55 min, 59 min, 60 min, 61 min and 65 min. The results are presented in Supplementary figure 16. Using a wrong *t_b_* had minimal effect on both *∂^max^* and V_ND_ estimates, and a big effect on *t_e_* and *t_s_* estimates.

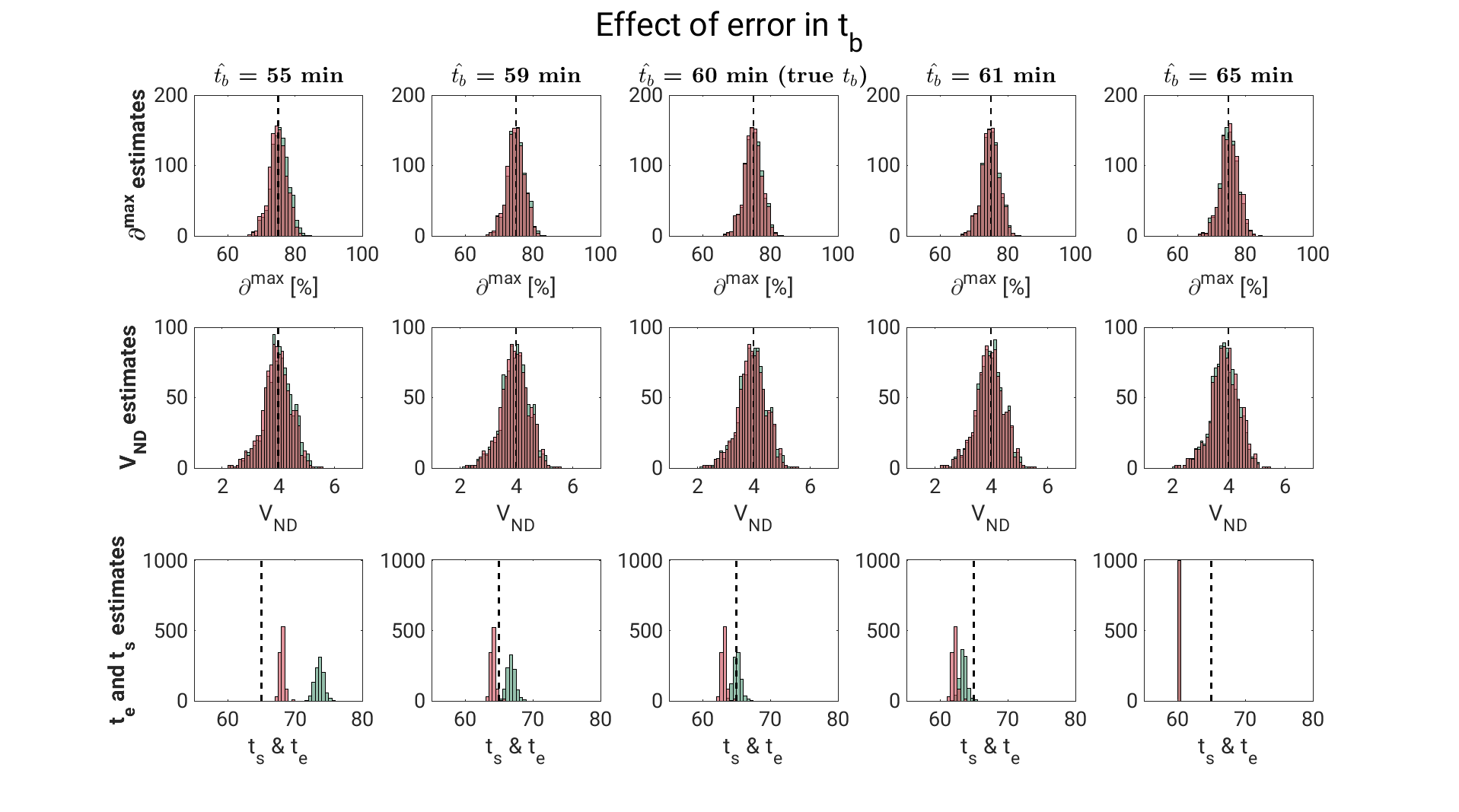

Supplementary figure 16: The effect of wrongly defined tb on parameter estimations. Histograms of ∂^max^, V_ND_, t_e_ (numerical solution) and t_s_ (single step solution) estimates for different values of t_b_. The true tb was 60 min. Estimates from the numerical solution are shown in green, and estimates from the single step solution are shown in red.
